# Antibiotics drive immunometabolic rewiring of human immune responses

**DOI:** 10.64898/2026.11.03.763842

**Authors:** Shelly Hen-Avivi, Noa Bossel Ben-Moshe, Ravit Avera, Inbal Adin, Yaarit Kitsberg, Shani K Lyskov, Shay Navon, Anat Zalmanovich, Tal Yardeni, Ronen Ben-Ami, Roi Avraham

## Abstract

Antibiotics are routinely evaluated for their antimicrobial activity, potential host toxicity, and disruption of the microbiome. Certain antibiotics can also impact cell-intrinsic host processes, including evolutionarily conserved processes in the mitochondria. These raise the possibility that antibiotics can directly impact host immunity, which relies on mitochondrial function for their immunometabolic activation. However, the immunological consequences of most antibiotics in current use have not been systematically examined. Here, we performed a comprehensive analysis of the effects of clinically used antibiotics on activation of primary human immune cells. We uncovered an underappreciated antibiotic-specific modulation of immune function in healthy individuals and infectious disease patients. Focusing on the immunomodulatory antibiotic ciprofloxacin, we found altered mitochondrial gene expression across multiple immune populations, driving distinct, lineage-specific responses and heightened sensitivity of T-cells. Ciprofloxacin impaired primary human T-cell expansion and activation and altered mitochondrial function and metabolism. We further demonstrated immunomodulatory effects of selected antibiotics in response to inactivated bacteria and during live viral infection. Finally, to test the implications of our findings *in vivo*, we showed that immune activation is altered shortly after a single dose of antibiotic administration in mice. This work established a widespread impact of antibiotics on immune-cell function, and identified mitochondrial perturbation as one underlying mechanism, with implications for host-pathogen interactions and antimicrobial therapy.

## Introduction

Antibiotics are critical for the treatment of bacterial infections and remain one of the most successful interventions in medicine. From the early pioneering work of Louis Pasteur and Paul Ehrlich^1^ to present-day research^2^, it is widely recognized that effective clearance of bacterial infections requires cooperation between antibiotics and the host immune system. The wide array of antibiotics in clinical use today, targeting various essential bacterial processes, rely heavily on immune-mediated clearance to achieve full resolution of infection. Bacteriostatic antibiotics, which inhibit bacterial growth (e.g., tetracyclines, macrolides), require the immune system to eliminate the impaired organisms^2^. Even bactericidal antibiotics, which actively kill bacteria in culture (e.g., fluoroquinolones, β-lactams), require the assistance of host immunity to clear residual bacteria. Furthermore, antibiotic concentrations at the site of infection may fall below minimal inhibitory concentrations (MIC) thresholds due to poor tissue penetration^3^, bacteria residing intracellularly^4^ or within biofilms^5^, underscoring the necessity of the immune system for bacterial clearance and treatment success^6,7^. This dependency is demonstrated in neutropenic mice, where antibiotic efficacy is significantly diminished, even for drugs that are highly effective in immunocompetent animals^8^. In cases of immunodeficient patients that are treated aggressively, treatment is hampered by the lack of a supportive immune system^9^.

Paradoxically, although considered to inhibit essential prokaryotic processes that are absent in eukaryotic cells, several antibiotics can induce host toxicity, including hepatic and renal dysfunction^10^, and disruption of the microbiome^11^. These processes can, in turn, indirectly modify host immune processes^11,12^. A more immediate, cell-intrinsic, and direct effect of antibiotics is their possible off-target effects on host cells, particularly the mitochondria, reflecting their evolutionary origin from α-proteobacteri^13^. These include mostly inhibition of translation^14^, inhibition of mitochondrial function^14,15^, and changes in metabolic pathways^16^. For example, aminoglycosides, which act by binding the 30S subunit of the bacterial ribosomal RNA, were shown to bind mitochondrial ribosomal RNA^17,18^. Chloramphenicol reversibly binds to the 50S subunit of the 70S ribosome in both prokaryotic organisms and mitochondria^19^. Ciprofloxacin inhibits bacterial DNA gyrase and topoisomerase IV, but can also interfere with mitochondrial topoisomerase II at higher concentrations^20^, thereby inhibiting metabolic activity of host cells^16^. These studies raise important questions about how widespread direct host immunomodulation is among antibiotics and the extent to which they may impair immune–antibiotic cooperation that is required for efficient bacterial clearance.

Previous studies indicate effects of individual antibiotics on immune phenotypes. For example, chlortetracycline markedly diminished phagocytosis of *Staphylococcus* by human leukocytes in vitro^21^. Macrolides have been demonstrated already in the 1970s to modulate inflammation and immunity, independent of antimicrobial properties^22^. Azithromycin reduces TNF-α mRNA levels and decreased TNF-α secretion, through NF-κB and specificity protein 1 (Sp1) DNA-binding activities^23^. Azithromycin was also shown to inhibit IL-1 secretion and non-canonical inflammasome activation^24^. Tetracycline was shown to significantly affect core pathways that modify host immunity^25,26^. Doxycycline was shown to confer protection in a mouse model of bacterial sepsis without affecting pathogen load^27^. Recent studies further indicate that immunomodulatory effects of antibiotics can impact infection outcome. For example, neomycin was shown to enhance host resistance to several viral pathogens^28,29^. Another study showed that bedaquiline reprogram macrophages into potent bactericidal phagocytes, and enhance intracellular killing of *Mycobacterium tuberculosis* strain that is insensitive to bedaquiline^30^. An additional layer of complexity is evident in studies suggesting both pro-inflammatory and anti-inflammatory effects, dependent on context, dose, infection status, and experimental model. For example, ciprofloxacin is mostly known to be immunosuppressive^31^, but also pro-inflammatory effects were shown^32^. Similarly, colistin was shown to have both pro- and anti-inflammatory effects^33,34^. These varying results indicate that the immunomodulatory effects may be context dependent and cell type-specific, requiring a better understanding of the underlying mechanism for antibiotic-induced immunomodulation.

It is becoming increasingly understood that immune cell activation is driven by rewiring of mitochondrial metabolism^35^. Mitochondria are now recognized as central regulators of immune cell fate decisions and antimicrobial function^36^, through immunometabolic reprogramming^37^. Detection of bacterial or viral ligands shift cellular metabolism, a process referred to as immunometabolic reprogramming^37^. This transition is actively orchestrated by mitochondria, with immune cell type-specific modifications. For example, a shift from OXPHOS to high-rate aerobic glycolysis in macrophages is driven by disruption of the tricarboxylic acid (TCA) cycle. Metabolic breakpoints reroute carbon flux and lead to accumulation of key intermediates such as succinate, citrate, and itaconate, which function as immunoregulatory signals^38^. The reliance of immune cells on mitochondrial processes suggests that antibiotics-mediated effects on mitochondrial processes may have profound implications on immunity during infection. Currently, most antibiotics in clinical use have not been systematically evaluated for their immunomodulatory potential, and even for previously identified immunomodulatory agents, a comprehensive mechanistic framework remains to be elucidated, representing a fundamental blind spot in antimicrobial therapy.

Here, we established an *ex vivo* screening platform based on isolated human peripheral blood mononuclear cells (PBMCs) from healthy donors. We systematically analyzed the effect of fifty clinically used antimicrobial drugs on immune activation in response to different ligands by measuring cytokine secretion, independent of their antimicrobial activity. To account for inter-individual variability, we selected eight immunomodulatory hits based on differences in cytokine profiles and analyzed their effects in healthy individuals as well as infectious disease patients. We performed a dose-response analysis of PBMCs incubated with increasing concentrations of antibiotic hits and measured dose-dependent immunomodulatory effect of our hits. Focusing on ciprofloxacin, we integrated single-cell transcriptomics, metabolic flux assays, and cell-cycle kinetics to establish that ciprofloxacin altered primary human T cells expansion and activation through disruption of mitochondrial processes. Finally, we demonstrate the implications of our findings *in vivo* in mice, where immune activation is altered shortly after a single dose of antibiotic administration. This work demonstrated that antibiotics may exert immunomodulatory effects that are highly context-dependent, shaped by immune cell identity, activation state, and mitochondrial processes.

## Results

### A comprehensive screen identified immunomodulatory effects of antibiotics in human PBMCs

We optimized conditions and readouts for screening human PBMCs using ligands that activate distinct immune programs. We extracted 5*10^8^ PBMCs from 500ml of blood from one donor and froze down in aliquots for each condition of the screen. To optimize ligand concentrations for detection of both increased and decreased cytokine secretion, PBMCs were incubated with varying concentrations of a toll-like receptor (TLR)-4 ligand lipopolysaccharide (LPS; 0.05–300 pg/ml), representing immune responses to Gram-negative infection, and a TLR3 ligand polyinosinic-polycytidylic acid (poly(I:C); 0.4–100 μg/ml), representing responses to viral infection. As immunological readouts, we quantified 12 cytokines and signaling molecules using commercially available HEK-Blue reporter cell lines, which detect TNFα, IL-17, IL-18, IFNγ, CD40L, IL-1R (detecting both IL-1α and IL-1β), IL-6, IFNα/β (detecting both IFNα and IFNβ), IL-2, IL-12, TGFβ, and IL-4/IL-13 (detecting both IL4 and IL13) (**Supplementary table S1**). Upon cytokine binding, the reporter cells secrete alkaline phosphatase into the culture medium, which was quantified using a colorimetric detection reagent and measured by spectrophotometry (**Supplementary figs. 1A, B**). Based on the dose-responses, we selected 50 pg/ml LPS or 50 μg/ml poly(I:C) in all subsequent experiments.

To systematically assess the immunomodulatory effects of clinically used drugs on human PBMCs, we screened a panel of 53 compounds comprising antibiotics from multiple classes, as well as antiviral, antifungal, and antiparasitic agents (**Supplementary table 2**). Dexamethasone, an anti-inflammatory drug, and cimetidine, a histamine H2 receptor antagonist, that are known immunomodulators were used as positive controls. To capture distinct immune activation programs, PBMCs were analyzed under three conditions: naïve (unstimulated), LPS-stimulated, or poly(I:C)-stimulated. The use of defined immune stimulants rather than live pathogens enabled assessment of drug-immune cell interactions independent of antimicrobial activity, thereby excluding indirect effects mediated through pathogen susceptibility. Antimicrobial drugs were added 1 h after stimulation and supernatants were harvested after 4 hours (h) and 24h for measurements of secretion levels of 12 cytokines. Baseline cytokine secretion in naïve PBMCs was low across most cytokines, whereas stimulation with LPS or poly(I:C) induced robust and stimulus-specific cytokine secretion (**Supplementary figs. 1C, D**). LPS stimulation predominantly induced pro-inflammatory mediators, including IFNγ, CD40L, and IL1R, whereas poly(I:C) stimulation enhanced a different profile of cytokines such as TNFα, IL-17, and IFNα/β. These distinct cytokine profiles confirm effective and differential immune activation by the two stimuli, with antibiotic-specific changes in cytokine secretion (**Supplementary figs. 1C, D**). Each antimicrobial was tested after 4h and 24h under all three conditions-naïve, LPS-stimulated, and poly(I:C)-stimulated PBMCs, and cytokine levels were quantified relative to a matched vehicle (no drug) control. We observed marked, time-dependent, and condition-dependent alterations in cytokine secretion by different antimicrobial drugs both after 4h and 24h (**Fig. 1, Supplementary fig. 1E**). To corroborate these results, we compared cytokine measurements by HEK-blue cells to Enzyme-Linked Immunosorbent Assay (ELISA) and measured similar cytokine changes induced by antibiotics (**Supplementary fig. 1F**). Noteworthy, immunomodulatory effects of individual antimicrobials were observed in one or two activation conditions, but not across all three tested states. For example, amikacin modulated cytokine secretion under naïve and poly(I:C)-stimulated conditions, whereas ciprofloxacin exerted immunomodulatory effects specifically under LPS- and poly(I:C)-stimulated conditions **(Fig. 1)**. These findings indicate that immunomodulatory activity is strongly dependent on the activation state of immune cells. Several antimicrobials attenuated stimulus-induced cytokine secretion, as observed for ciprofloxacin and levofloxacin under both LPS and poly(I:C) stimulation **(Fig. 1)**. In contrast, other compounds enhanced specific cytokine responses, including amikacin, neomycin, and kanamycin under poly(I:C) stimulation, highlighting the diversity of immunomodulatory activity across antimicrobials. In some cases, we observed similar effects for antibiotics from the same class: aminoglycosides broadly enhanced responses to viral ligands, and fluoroquinolones suppressed responses to Gram-negative bacterial stimuli. Finally, antibiotics previously reported to exhibit immunomodulatory properties, such as neomycin^28,29^ and bedaquiline^30^, were identified as hits in the screen, providing internal validation of the experimental approach.

**Figure 1:**
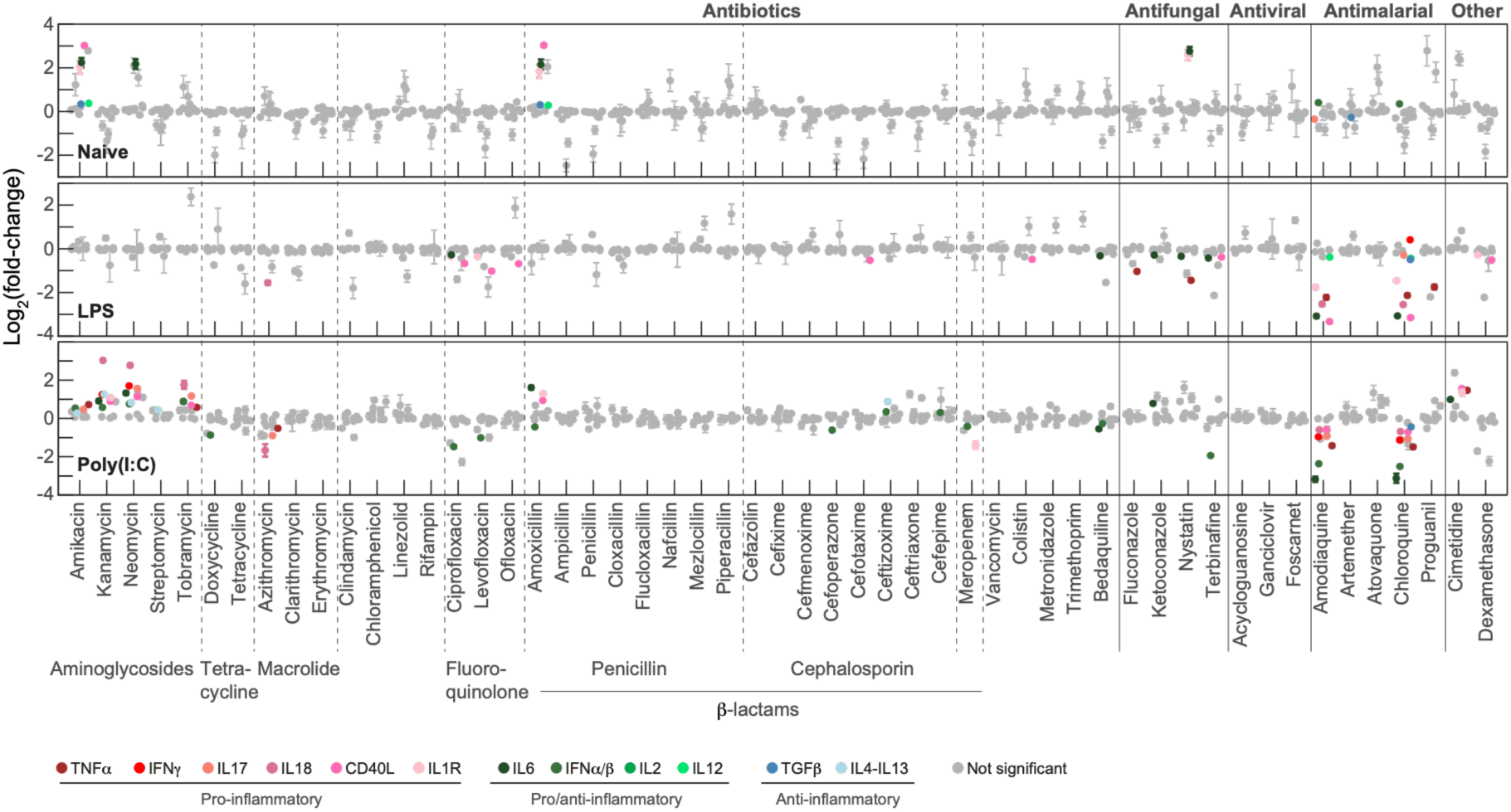
A comprehensive screen identified immunomodulatory effects of antimicrobial drugs. PBMCs from a healthy donor were either unstimulated (upper) or stimulated with LPS (middle) or poly(I:C) (lower), exposed to the indicated drugs (X-axis, antibiotic concentrations in Supplementary table 2) for 24h, and tested for secretion of a panel of 12 cytokines (indicated in the colorbar). For each condition, log2(fold change) between with and without drug was calculated (Y-axis). Non-significant changes are shown in grey, and significant changes with 1%FDR are colored (2-sample t-test). Results are from four replicates presented as mean ± standard error.

### Inter-individual variability in the immunomodulatory effect of antibiotics

We selected eight antibiotics identified as immunomodulatory hits in the primary screen (amikacin, neomycin, azithromycin, amoxicillin, meropenem, colistin, ciprofloxacin, levofloxacin) for further analyses. First, we evaluated potential cytotoxicity and measured no significant changes in PBMC viability following 24 h of treatment in the concentrations tested, indicating that the observed changes in cytokine secretion were not attributable to cellular toxicity (**Supplementary fig. 2A**). Next, to assess inter-individual variability in drug-mediated immune modulation, PBMCs were obtained from a cohort of ten healthy donors. We analyzed the immunomodulatory effects of the eight antibiotic hits under naïve, LPS-stimulation, or poly(I:C)-stimulation and measured cytokine secretion. Dimensionality reduction using t-distributed Stochastic Neighbor Embedding (t-SNE) of cytokine secretion profiles revealed a clear segregation of samples primarily driven by stimulation condition, suggesting distinct immune activation programs (**Fig. 2A, Supplementary fig.2B**). Within each condition, we observed substantial inter-individual variation in baseline immune responses as well as in responses to stimulation and antibiotic treatment, reflected by the wide distribution of the responsive space (dashed circles in **Fig. 2A**, dashed regions occupied for each stimulant without antibiotic treatments are marked for visualization only). Several antibiotics induced consistent shifts in sample localization away from the no antibiotic reference regions across all individuals (*e.g.,* neomycin and ciprofloxacin), whereas others affected only a subset of individuals (*e.g.,* Amoxicillin and colistin), indicating both antibiotic-specific and individual-specific immunomodulatory effects. To quantify these effects, we stratified the differences between individuals in cytokine secretion to each antibiotic. We revealed antibiotics with high variation in their immunomodulatory activity (*e.g*., amoxicillin) and antibiotics that are immunomodulatory across all individuals tested (*e.g.,* ciprofloxacin) (**Fig. 2B**).

**Figure 2:**
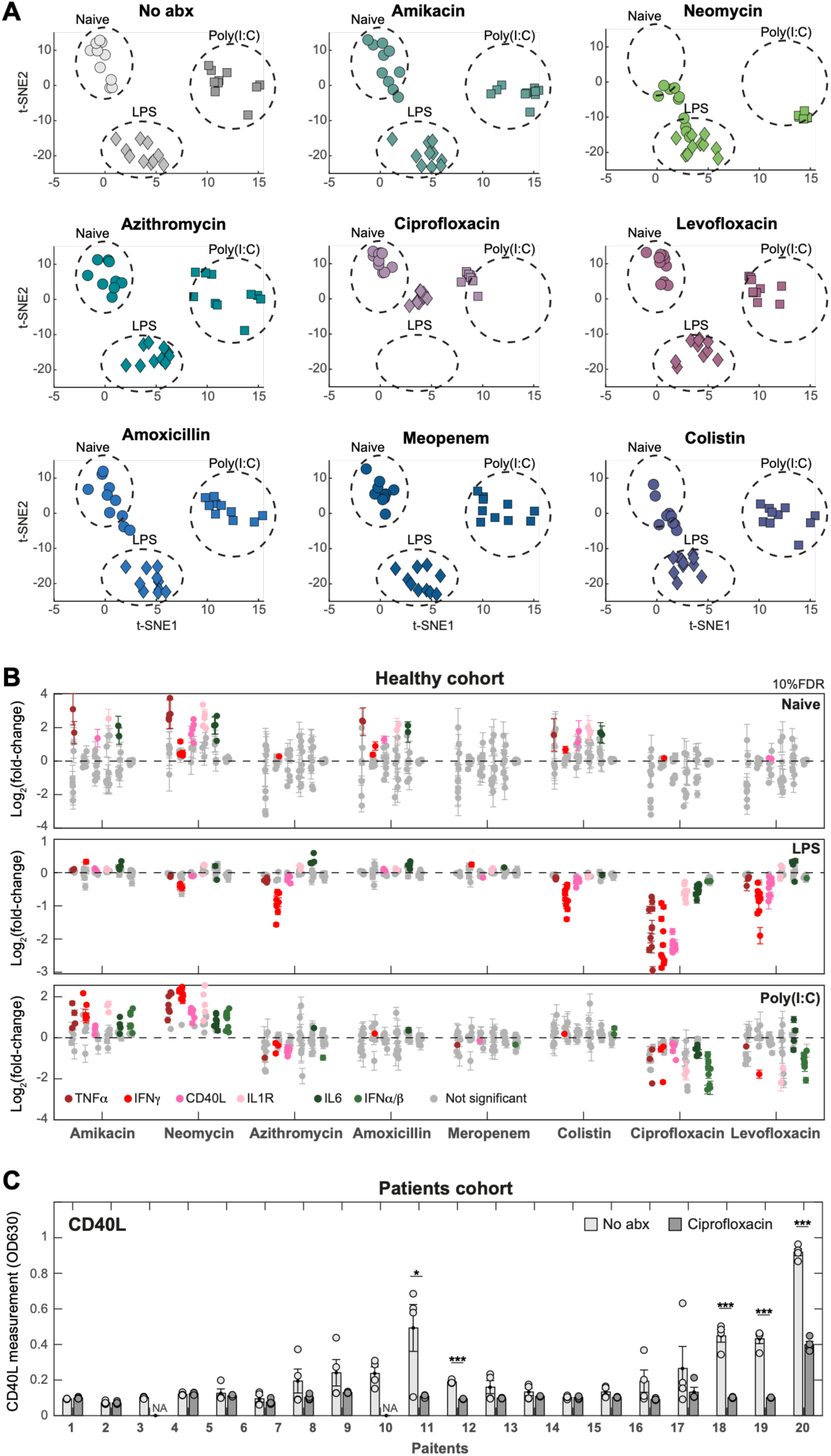
Analyzing inter-individual variation of the effects of immunomodulatory antibiotics. (**A+B**) PBMCs from ten healthy individuals were either unstimulated or stimulated with LPS or poly(I:C), exposed to the indicated drugs for 24h (no abx-no antibiotics control). (**A**) tSNE representation of cytokine secretion measurements for each drug. Dotted circles mark the area each stimulant occupies in no antibiotic control condition. (**B**) For each antibiotic (x-axis), each column presented is a different cytokine measurement (indicated in the colorbar, non-significant changes are shown in grey), and each dot in the column is an individual. For each condition, log2(fold change) between with and without drug was calculated (Y-axis) (2-sample t-test, 10%FDR). (**C**) PBMCs from 20 patients (X-axis) were collected and exposed to ciprofloxacin for 24h and tested for CD40L levels (Y-axis). *P* values indicated are from 2-sample t-test, * p<0.05, ** p<0.01, *** p<0.001.

To test our findings in a more clinically relevant setting, we collected PBMCs from a cohort of 20 hospitalized patients with diverse clinical backgrounds (**Supplementary table S3**). Patient-derived PBMCs were either left untreated or treated with the immunomodulatory hit ciprofloxacin (**Fig. 2C**, and **Supplementary figs. 2C-D**) or amikacin (**Supplementary fig. 2E**). We measured a significant reduction in cytokine levels upon ciprofloxacin treatment only in PBMCs samples that displayed elevated baseline cytokine secretion, probably due to their inflammatory disease-related state. For amikacin, the immunomodulatory effects were more variable, mirroring the variation observed also in healthy individuals. These immunomodulatory effects were not correlated with patients’ age or sex (**Supplementary fig. 2F**). These results indicate that immunomodulatory antibiotics identified in our screen may have effects beyond inter-individual variation and in patients with active inflammation.

### Dose-dependent and activation state-dependent immunomodulatory effects of antibiotics

To further support the specificity of the immunomodulatory effects induced by antibiotic hits, we assessed their dose-dependent activity on immunity. PBMCs were exposed to increasing concentrations of selected hit antibiotics under LPS or poly(I:C) stimulation, and cytokine secretion was measured. We observed a clear, dose-dependent modulation of immune responses (*e.g.,* CD40L in LPS-stimulated samples and IFNα/β in poly(I:C)-stimulated samples; **Fig. 3A, Supplementary fig. 3A**). These may be relevant to the clinical setting, as the dose-dependent immunomodulatory effects could be observed at plasma *C*max (**Supplementary table 4**), which reflects the concentrations to which circulating PBMCs may be exposed during antibiotics treatment. We next performed a set of experiments to test the possibility that antibiotic immunomodulation is specific to immune activation states. PBMCs were subjected to different sequences of stimulation and antibiotic exposure (amikacin and ciprofloxacin): cells were either stimulated for 1 h followed by addition of an immunomodulatory antibiotic or treated first with the antibiotic for 1 h prior to stimulation. We measured cytokine secretion after 24 h and observed that both antibiotics had immunomodulatory effects when added either before or after stimulation (LPS with CD40L and IL6 for ciprofloxacin, and poly-(I:C) with IFNα/β and IFNγ for amikacin, **Fig. 3B, Supplementary fig. 3B**). We then performed a similar experiment, but adding a wash step: cells were stimulated for 1 h, the stimulant was washed out, and the antibiotic was subsequently added. Under these conditions, antibiotics retained their immunomodulatory activity. In contrast, when cells were pretreated with the drug for 1 h followed by a wash of the antibiotics and subsequent addition of the stimulant, the immunomodulatory effect was completely abolished (**Fig. 3C, Supplementary fig. 3C**). These results suggest that antibiotic-mediated immunomodulation is context-dependent, and not simply general toxicity, with effects that target a specific immune activation cell state.

**Figure 3:**
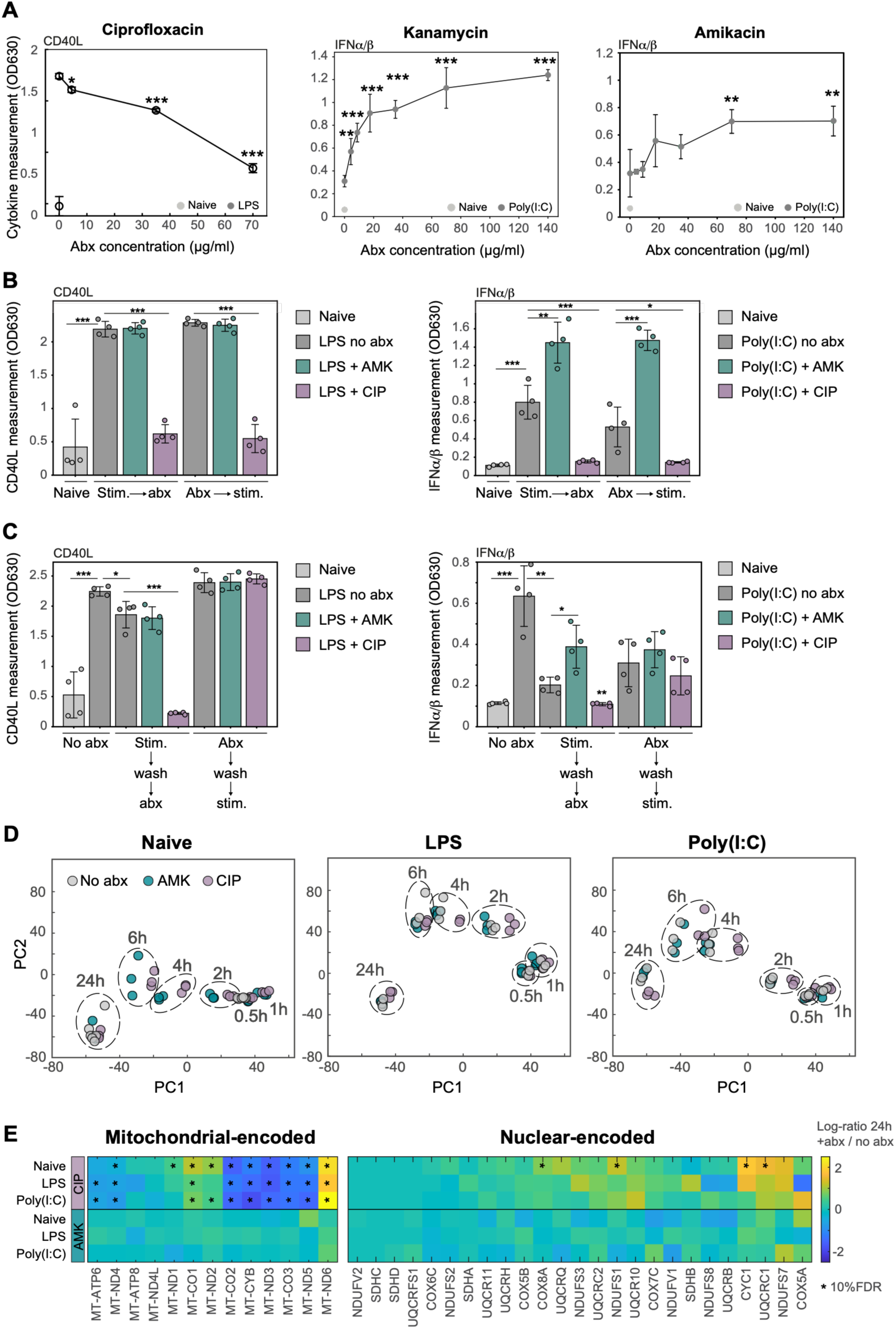
Dose-dependent and activation state-dependent immunomodulatory effects of antibiotics. (**A**) Indicated are the cytokine measurements of PBMCs from a healthy donor, either naïve (light grey) or stimulated (dark grey), and treated with different doses of the indicated antibiotics (X-axis). (**B**) PBMCs were either stimulated with LPS (left) or poly(I:C) (right) before or after antibiotics exposure (indicated on X-axis; AMK-amikacin, CIP-ciprofloxacin, stim-stimulant, abx-antibiotics) and measured for cytokine secretion (Y-axis). Naïve PBMCs are indicated in light grey and stimulated PBMCs without antibiotics in dark grey. (**C**) PBMCs were stimulated with LPS (left) or poly(I:C) (right), stimulant was washed after 1h and the indicated antibiotics were added (X-axis) and measured for cytokine secretion (Y-axis). Alternatively, PBMCs were treated with the indicated antibiotics for 1h, antibiotics were washed and stimulant was added. Naïve aBMCs are indicated in light grey and stimulated PBMCs without antibiotics in dark grey. (**D**) PBMCs from a healthy donor were kept unstimulated, or stimulated with LPS, or poly(I:C), in the presence or absence of amikacin and ciprofloxacin. At the indicated time points, the cells were subjected to bulk RNAseq analysis. PCA computed using the complete dataset across all experimental conditions combined, with individual plots displaying samples corresponding to specific stimulation states (naïve, LPS, or poly(I:C)) and either untreated or treated with ciprofloxacin (CIP) or amikacin (AMK) at different time points. (**E**) Heatmaps displaying fold changes at 24h between ciprofloxacin-or amikacin-treated samples and control (no antibiotics) of ETC complex genes, either mitochondrial encoded (left) or nuclear encoded (right) from bulk RNA seq data. * Indicates 10%FDR from 2-samples t-test comparing between no antibiotics and antibiotics treatment. *P* values indicated are from 2-sample t-test, * p<0.05, ** p<0.01, *** p<0.001. Antibiotic concentrations in B-E: amikacin-70 mg/ml, ciprofloxacin-70 mg/ml.

To test possible mechanisms of immunomodulatory effects of antibiotics, we performed a time course analysis by RNA sequencing (RNA-seq) of PBMCs, either unstimulated, or stimulated with LPS, or poly(I:C), in the presence or absence of amikacin and ciprofloxacin. We examined principal component analysis (PCA) of all samples and observed that PC1 predominantly captured temporal effects, whereas PC2 captured stimulation-associated variation (**Fig. 3D**). PC1 pattern indicated that time-dependent transcriptional changes occurring during 24 h were also evident in resting cells, suggesting these are culture related changes. In stimulated samples, we observed that amikacin induced transcriptional changes mostly at early time points compared to untreated samples. Ciprofloxacin induced pronounced transcriptional changes throughout the time course and persisted through the 24 h time point under both LPS and poly(I:C) stimulation. We then performed differential gene expression analysis across all conditions and identified sets of genes for each antibiotic, and enriched immune pathways associated with ciprofloxacin treatment only (**Supplementary figs. 4A-C**). To test our hypothesis for the involvement of mitochondrial processes in antibiotic-induced immunomodulation, we specifically examined mitochondrial gene expression of mitochondrial-encoded mRNAs and nuclear-encoded mRNAs that are imported into mitochondria (**Fig. 3E**). Interestingly, ciprofloxacin inhibited the expression of most mitochondrial-encoded genes, suggesting an effect on mitochondrial function. In contrast, one mitochondrial-encoded gene, complex I subunit ND6, was induced by ciprofloxacin. Notably, ND6 is the only protein encoded on the light strand of the mtDNA and is less susceptible to stress during mtDNA replication^39^. These results prompted us to directly examine the relationship between mitochondrial function and cytokine production. We measured cytokine production from PBMCs that were stimulated with LPS or LPS in the absence or presence of rotenone, an inhibitor of mitochondrial electron transport chain complex I. Direct inhibition of mitochondrial respiration with rotenone resulted in a marked reduction in TNF-α and IFN-γ secretion compared with LPS stimulation alone (**Supplementary figs. 4D-E**). These findings demonstrate that intact mitochondrial function is required to sustain the cytokine responses measured in our PBMCs experimental system.

### Transcriptional analysis indicated shared as well as cell type-specific mitochondrial gene expression alterations by ciprofloxacin

To analyze cell type-specific responses to the immunomodulatory antibiotic ciprofloxacin, we performed single-cell RNA-seq (scRNA-seq) on resting and LPS-stimulated PBMCs from 2 healthy donors, either untreated or treated with ciprofloxacin. We tested ciprofloxacin across a range of concentrations (13.5, 105, or 210 μM), with the lowest concentration corresponding to the reported human plasma *C*max (13.5 µM; **Supplementary table 4**), based on human pharmacokinetic data following a 750-mg oral dose. Cells were clustered and classified based on canonical marker genes into natural killer (NK) cells, NKT cells, T cells, B cells, monocytes, macrophages, and dendritic cells. We further sub-clustered each major cell type to resolve the repertoire of immune-cell subsets (**Supplementary figs. 5A-B**). Two-dimensional t-distributed Stochastic Neighbor Embedding (t-SNE) visualization illustrated differences in the distribution of cellular states across donors, stimulation conditions, and ciprofloxacin treatment (**Fig. 4A**). The relative abundance of several immune-cell subsets changed in response to ciprofloxacin, in a dose-dependent manner (*e.g.,* cytotoxic CD8/γ/δ (6), memory CD4 T cells (9), activated CD4 memory T (21), activated/stressed T (4); **Supplementary fig. 5B**, bottom), also at low ciprofloxacin concentrations (**Supplementary fig. 5C**).

**Figure 4:**
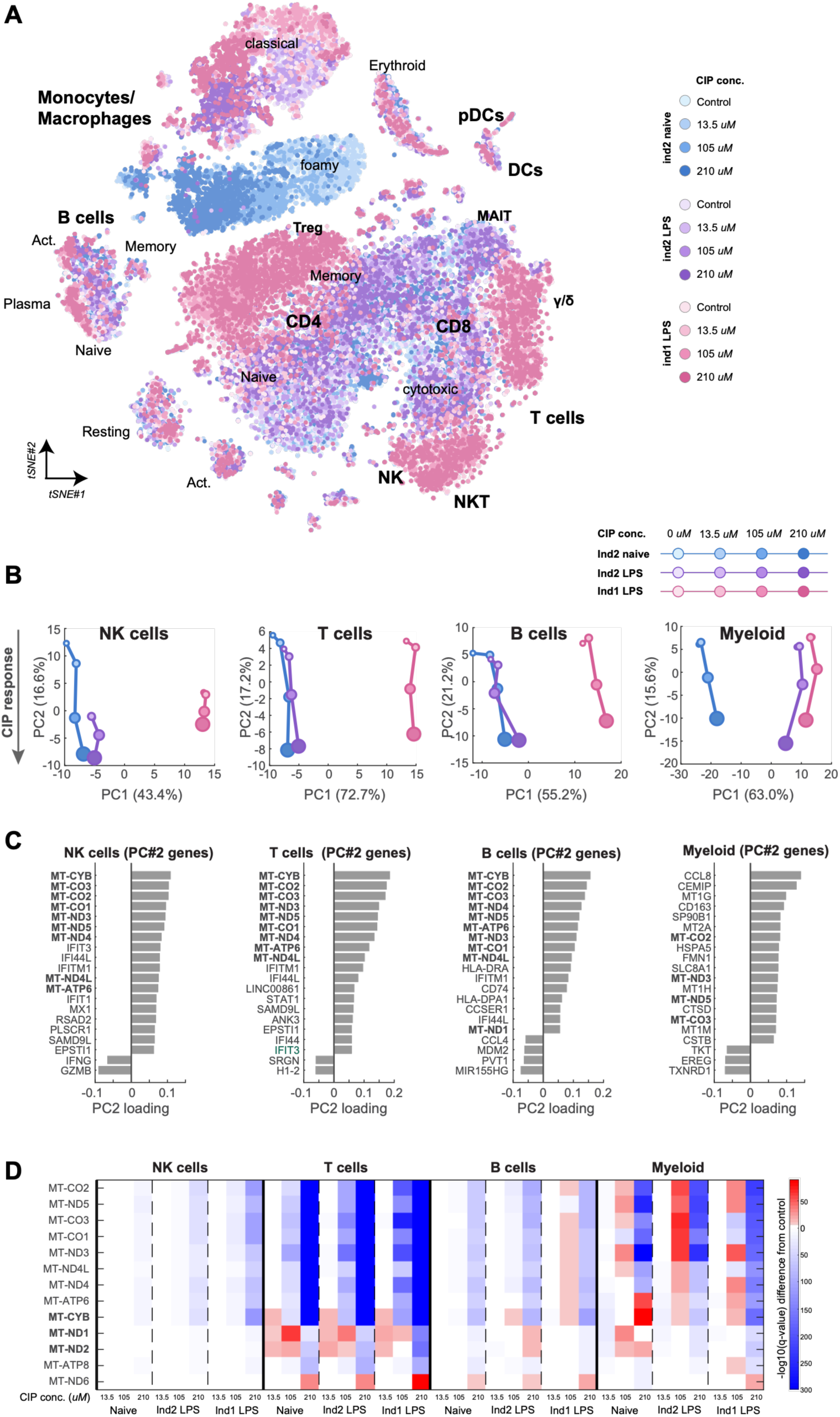
Transcriptional analysis indicated cell type-specific mitochondrial gene expression alterations by ciprofloxacin. PBMCs from two healthy donors were unstimulated or stimulated with LPS and untreated or treated with ciprofloxacin (13.5, 105, 210 μM) for 24h, and analyzed by scRNA-seq. (**A**) tSNE visualization of scRNA-seq data combining the two individuals and treatments together. Cell-type annotations and color legends are provided. (**B**) Principal component analysis (PCA) projection of the mean scRNA-seq data of each cell type, based on all expressed genes, onto the space of the two leading principal components for all samples together. (**C**) Dynamics of the percentage of cells across different ciprofloxacin concentrations per individual and stimulation within representative cell subsets. (**D**) Heatmap displaying -log10(q-value) of all mtDNA genes (Y-axis) in response to ciprofloxacin in different concentrations (X-axis) in the different individuals and stimulants per cell-type (colorbar indicates significant changes, with 10% FDR). *P* values are from 2-sample t-test. Highlighted genes are significant at 13.5 μM ciprofloxacin.

To assess cell-type-specific transcriptional responses to ciprofloxacin under unstimulated and LPS-stimulated conditions, we performed Principal Component Analysis (PCA) for each major cell type individually (NK cells, T cells, B cells, and myeloid cells). PC2 was associated with ciprofloxacin treatment in a dose-dependent manner across all cell types, donors, and stimulations (**Fig. 4B**). The transcriptional signatures associated with the ciprofloxacin response, represented by genes with high PC2 loadings, were strongly enriched for mitochondrial DNA (mtDNA)-encoded genes (**Fig. 4C**). Further examination of all mtDNA-encoded genes revealed differences in the magnitudes of response across cell populations, with T cells exhibited the most pronounced changes in mitochondrial transcripts following ciprofloxacin exposure under both unstimulated and LPS-stimulated conditions (**Fig. 4D** and **Supplementary fig. 5D**). Other immune-regulated genes (*e.g.,* TNIP2, a negative regulator of NF-kB signaling that suppresses the production of pro-inflammatory cytokines^40^ were also among the top upregulated genes in T cells following ciprofloxacin treatment (**Supplementary fig. 5E, Supplementary table 5**). Together, these scRNA-seq data provide cell-type-resolved evidence that ciprofloxacin induced transcriptional reprogramming across multiple immune-cell populations and identify T cells as particularly sensitive to its effects on mitochondrial gene expression.

### Ciprofloxacin impacted mitochondrial respiration of T-cell and inhibited proliferation

To test directly the possible effects of ciprofloxacin on the mitochondrial function and activation of T cells, we implemented a protocol of T cell expansion from human PBMCs. T cells were activated and expanded using soluble anti-CD3 and anti-CD28 antibodies to mimic T cell receptor (TCR) engagement and co-stimulatory signaling, together with IL-2 to support T cell proliferation^41,42^. We conducted RNA-seq on activated T cells treated with three concentrations of ciprofloxacin. PCA analysis revealed dose-dependent transcriptional changes induced by ciprofloxacin (**Fig. 5A**), observed on PC1 genes, including mitochondrial genes (**Fig. 5B**, **Supplementary fig. 6A**). We analyzed DEGs between untreated and ciprofloxacin treated samples (**Supplementary fig. 6B**), and measured enrichment of pathways (**Supplementary fig. 6C**), including mitochondrial metabolism and T cell proliferation (**Fig. 5C**). These effects of ciprofloxacin on mitochondrial genes, corroborated our scRNA-seq results, and prompted us to us to test directly the effects of ciprofloxacin on mitochondrial metabolism and proliferation of T cells.

**Figure 5:**
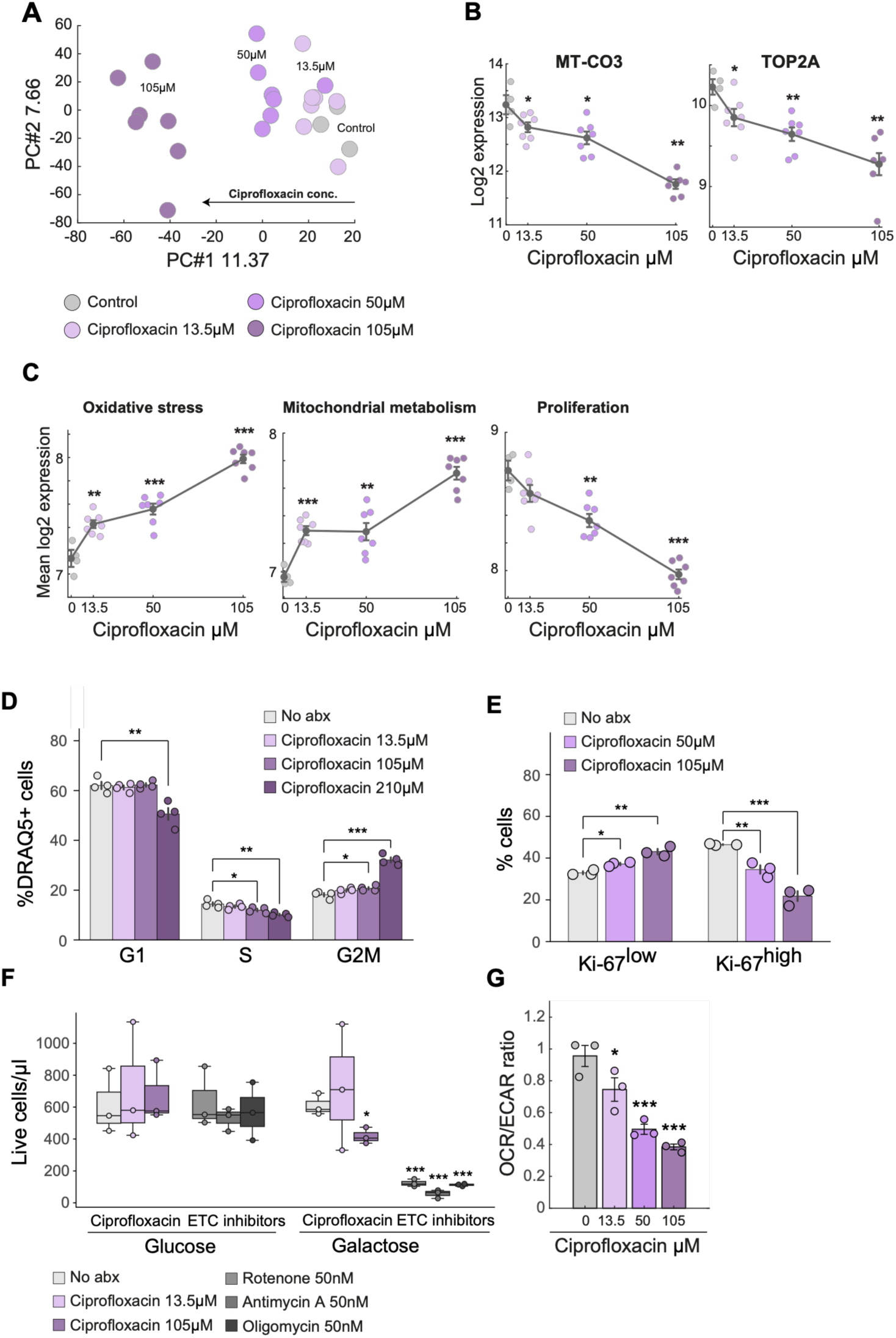
Ciprofloxacin impacted mitochondrial respiration of T-cell and inhibited proliferation. (**A-C**) T cells were activated from PBMCs, treated with the indicated concentrations of ciprofloxacin for 4 days and subjected to bulk RNA-seq. **(A)** Principal component analysis (PCA) projection of the RNA-seq data based on all expressed genes onto the space of the two leading principal components for all samples together. PC1 demonstrates a distinct dose-dependent trajectory with increasing ciprofloxacin concentration (indicated by the arrow). **(B)** Log_2_ expression levels (Y-axis) of representative PC1-associated genes, MT-CO3 and TOP2A, across ciprofloxacin treatment concentrations (X-axis). **(C)** Mean log_2_ expression levels (Y-axis) of representative biological pathways that are changed across ciprofloxacin doses (X-axis). (**D**) T cells were treated with the indicated concentrations of ciprofloxacin for 24h, stained with an antibody against Ki-67 and with DRAQ5 DNA-binding dye, and cell cycle was analyzed using flow cytometry. Indicated are the quantification of cell cycle phases based on DRAQ5 from Ki-67^high^ population. (**E**) T cells were treated with the indicated concentrations of ciprofloxacin for four days and proliferating cells were quantified with an antibody against Ki-67. (**F**) Quantification of T cell viability in glucose vs. galactose media under ciprofloxacin treatment, measured by flow cytometry. Mitochondrial ETC inhibitors (Rotenone, Antimycin A or Oligomycin) were used as controls. Bar plots represent the absolute number of live cells per sample as normalized by counting beads. Live/dead cells were determined by Sytox-blue staining. (**G**) T cells from three healthy donors were treated with the indicated concentrations of ciprofloxacin or control for 4 days. On day five, the ratio of oxygen consumption rate (OCR) to extracellular acidification rate (ECAR) was measured using a Seahorse XF extracellular flux analyzer. Bars represent mean ± SEM of 3-6 technical replicates. *P* values indicated are from ANOVA and Dunnett’s multiple comparison test, compared to the untreated control (no antibiotics). *P* values indicated are from 2-sample t-test, * p<0.05, ** p<0.01, *** p<0.001.

To test the effects of ciprofloxacin on T cell proliferation we analyzed by flow cytometry Ki-67, a proliferation marker, and DRAQ5, a DNA-binding dye used to analyze cell-cycle progression. We validated T cell expansion and activation by flow cytometry using CD134 as an activation marker for CD4⁺ T cells and CD137 for CD8⁺ T cells^43^. Both activation markers were robustly induced four days following CD3/CD28 stimulation, and declined by day seven (**Supplementary figs. 7A, B**). Next, T cells were treated with ciprofloxacin at different concentrations for 24 h. Using flow cytometry, we gated on Ki-67^high^ (proliferating) and Ki-67^low^ (non-proliferating) populations (**Supplementary fig. 7C**). Within the Ki-67^high^ proliferating cells we used DRAQ5, a cell-permeable fluorescent DNA-binding dye that enables quantification of cellular DNA content, to determine the proportions of cells in the G1, S, and G2/M phases of the cell cycle (**Supplementary fig. 7D**). We observed a significant and dose-dependent effect of ciprofloxacin on the proportion of cells in S and G2/M phase that were gated from the Ki-67^high^ population (**Fig. 5D**). Next, we assessed a longer course of four days ciprofloxacin treatment on T cells. We measured a dose-dependent decrease in Ki-67^high^ proliferating cells and an increase in Ki-67^low^ populations (**Fig. 5E, Supplementary fig.7E**). Together, these findings indicate that ciprofloxacin affects cell cycle of T cell and their proliferation during activation.

We next turned to test whether the effects of ciprofloxacin on mitochondrial genes and T cell proliferation may be associated with mitochondrial dysfunction. For this, we compared ciprofloxacin-treated T cells cultured in galactose vs. glucose media. As galactose is not efficiently incorporated into glycolysis, cells are forced to rely primarily on mitochondrial oxidative phosphorylation for ATP production. Cells cultured in glucose can compensate for mitochondrial defects through glycolysis. Consequently, compounds that impair mitochondrial function selectively reduce cell viability under galactose conditions^44^. T cells were treated with ciprofloxacin or PBS for 4 days, and then the culture media were replaced with either glucose- or galactose-containing media for 24 h, in the presence or absence of ciprofloxacin. Electron transport chain (ETC) inhibitors (rotenone; complex I, antimycin A; complex III, and oligomycin; complex V) were included as positive controls. We observed that ciprofloxacin induced significant cell death in galactose-containing media, but not in glucose-containing media (**Fig. 5F**). These effects were correlated with the effects of ETC inhibitors, indicating that ciprofloxacin disrupts mitochondrial function, linking its effects on T cell proliferation to impaired mitochondrial respiration.

Finally, we studied the direct effects of ciprofloxacin on mitochondrial respiration in T cells. We expanded T cells from three healthy donors and treated the cells with ciprofloxacin for 4 days at three concentrations. On day five, we performed Seahorse mito stress test, and measured oxygen consumption rate (OCR; **Supplementary fig. S8A**) and extracellular acidification rate (ECAR; **Supplementary fig. 8B**). Ciprofloxacin reduced significantly OCR/ECAR ratios in all three donors, indicating a shift in cellular respiratory balance and reduced mitochondrial respiratory activity (**Fig. 5G, Supplementary fig. 8C**). In parallel, we performed quantitative PCR measurements of mitochondrial DNA (mtDNA) in the same three donors and measured effects of ciprofloxacin on mitochondrial content only in one individual (**Supplementary fig. 8D-E**). These results suggest that ciprofloxacin induces changes to mitochondrial respiration of T cells, irrespective of mitochondrial numbers.

### Immunomodulatory antibiotics impacted cytokine secretion *in vivo* and during infection

To evaluate host-directed immunomodulation of antibiotics in response to bacterial challenge, irrespective of their direct antimicrobial activity, PBMCs from a healthy donor were stimulated with heat-killed *E. coli* across a range of ciprofloxacin concentrations (13.5, 52.5, 105, and 210 μM). Consistent with observations in LPS-stimulated cells, heat-killed *E. coli* robustly induced the secretion of CD40L, IL6, and TNFα. Ciprofloxacin treatment significantly suppressed this response (**Fig. 6A**). To further test the effects of immunomodulatory antibiotics in response to a live pathogen, we infected PBMCs from three healthy donors with human cytomegalovirus (HCMV) TB40 strain. As antibiotics lack direct antiviral activity, this model enabled us to examine host-directed immunomodulatory effects during live infection without confounding by direct antimicrobial activity against the pathogen. The HCMV carries triple fluorescent tags: mNeoGreen for immediate early gene, mTagBFP2 for early gene, and mCherry for late viral gene expression, enabling temporal validation of viral infection^45^. We followed the kinetics of viral infection and replication within infected cells based on the fluorescence signal of HCMV. Using flow cytometry, we were able to detect infection with HCMV by 18 h post-infection (**Supplementary fig. 9A**). Kanamycin, amikacin, colistin, azithromycin and ciprofloxacin were selected for viral infection experiments based on their immunomodulatory effects observed under poly(I:C) stimulation. HCMV infection of PBMCs resulted in a significant increase in IFNα/β and IFNγ secretion (**Supplementary fig. 9B)**. Notably, kanamycin and colistin increased IFNβ secretion (in 2 out of 3 individuals) and IFNγ secretion in one individual. Ciprofloxacin significantly decreased IFNβ in all individuals and IFNγ secretion in one individual. We measured significant effects of amikacin and azithromycin only in one individual (**Fig. 6B**). These findings indicate that immunomodulatory antibiotics which alter cytokine responses in stimulant-based assays extend their effects to the context of viral infection, with observed variation among different individuals (**Fig. 2B**).

**Figure 6:**
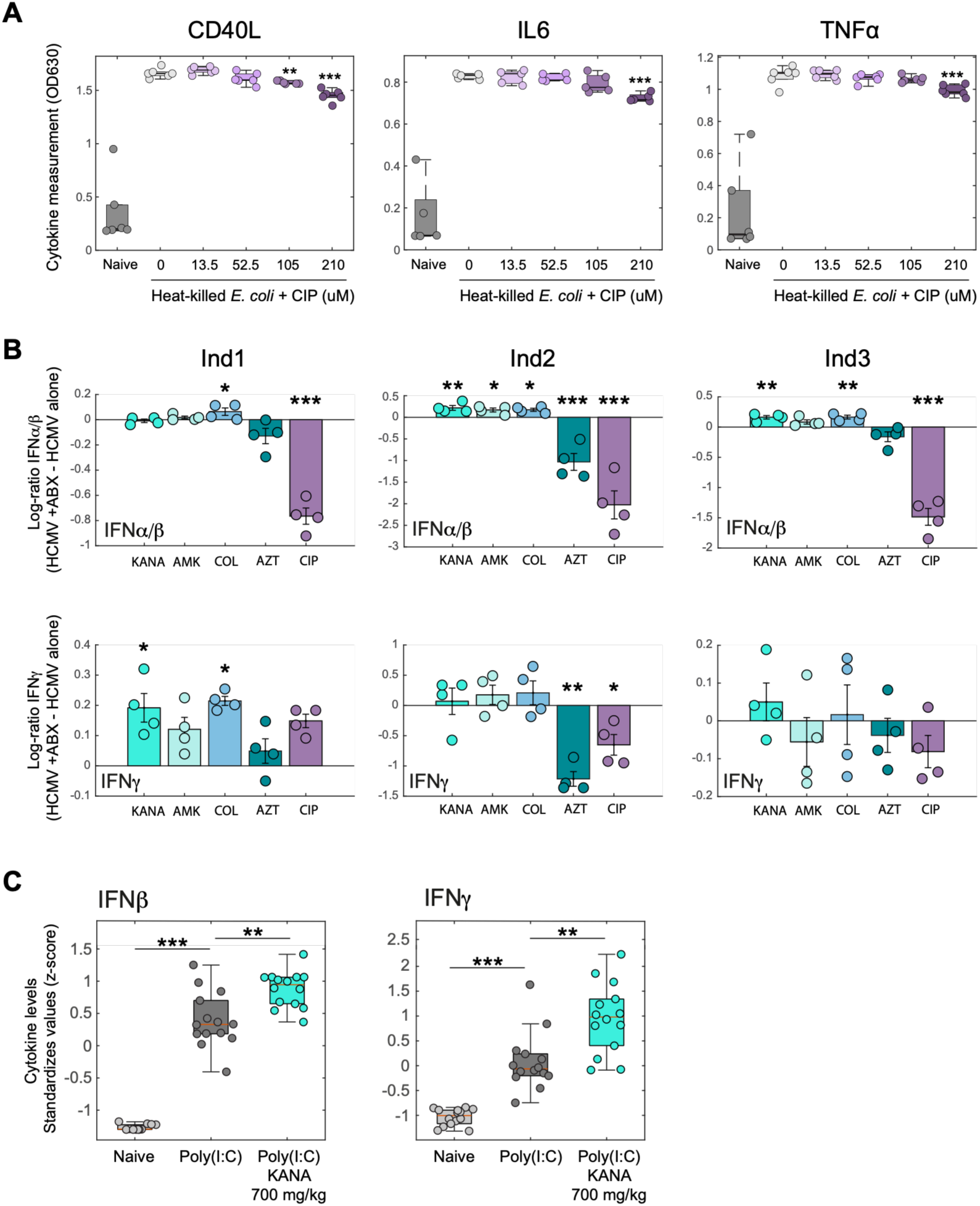
Immunomodulatory antibiotics impact cytokine secretion during infection. (**A**) PBMCs from a healthy donor were either untreated or treated with heat-inactivated *E. coli,* then exposed to different ciprofloxacin concentrations (0, 13.5, 52.5, 105 or 210 μM). Changes in CD40L, IL6 and TNFα were measured by HEK-blue reporter cell lines. (B) PBMCs from three healthy donors were infected with HCMV, treated with the indicated antibiotics or control. After 18 h IFNα/β (upper panel) and IFNγ (lower panel) cytokine secretion was measured. Y-axis indicates the log-ratio of cytokine secretion of antibiotic treatment vs. no antibiotic control, measured by HEK-blue reporter cell lines. KANA-kanamycin (40 μg/ml), AMK-amikacin (70 μg/ml), COL-colistin (3 μg/ml), AZT-azithromycin (100 μg/ml), CIP-ciprofloxacin (70 μg/ml). (**C**) Mice were injected intravenously with 150 μg/mouse poly(I:C), followed immediately by subcutaneous administration of kanamycin (700 μg/kg). Four hours post-injection, mice were sacrificed, and blood was collected by retro-orbital bleeding. Plasma was isolated and cytokine levels were quantified by ELISA for IFNβ (lest panel) and IFNγ (right panel), presented as Z-scores (Y-axis) from three independent experiments. Each dot represents measurements from a single animal. *P* values indicated are from 2-sample t-test, * p<0.05, ** p<0.01, *** p<0.001.

Next, we sought to evaluate the immunomodulatory activity of antibiotics in the *in vivo* setting using a mouse model. As immunometabolic processes are known to differ between mouse and human immune cells^46^, we tested the effects of several antibiotics we identified as immunomodulatory in human immune cells in mouse splenocytes. Splenocytes from naïve mice were isolated, stimulated *ex vivo* with ligands, treated with a panel of selected antibiotic hits, and measured cytokines using HEK reporter assays (**Supplementary fig. 9C**). Importantly, kanamycin elicited a strong immunomodulatory response in murine immune cells, similar to that we observed in human PBMCs, making it a suitable candidate for *in vivo* studies. However, ciprofloxacin showed relatively modest effects in murine immune cells, unlike its effects in human PBMCs. Next, we injected mice intravenously with poly(I:C) (150μg/mouse) followed immediately by subcutaneous administration of kanamycin (500 mg/kg, 700 mg/kg or 1100 mg/kg). Four hours post-injection, mice were sacrificed, and blood and spleens were collected. Plasma was isolated and cytokine levels were quantified by ELISA. Poly(I:C) injection induced a significant increase in IFNβ and IFNγ levels (**Fig. 6C** and **Supplementary fig. 9D**). Notably, kanamycin increased both IFNβ and IFNγ levels compared to mice treated with poly(I:C) only, indicating the immunomodulatory effects of kanamycin *in vivo*. Spleens were homogenized and analyzed by bulk RNAseq to test for the effects of kanamycin on gene expression. We measured poly(I:C)-induced transcriptional changes in mouse splenocytes (**Supplementary Fig. 9E** and **Supplementary Table 7).** Importantly, kanamycin treatment further increased an inflammatory granulocyte signature while decreasing a macrophage-associated signature (**Supplementary Fig. 9F**), suggesting that kanamycin affects immune cell composition and/or infiltration in vivo. Finally, we tested the immunomodulatory effects of kanamycin on downstream immune-cell function. We therefore examined macrophage phagocytic capacity using a fluorescent bead uptake assay. PMA-differentiated THP1 cells were exposed to conditioned supernatants from PBMCs stimulated with poly(I:C), in the presence or absence of kanamycin, followed by incubation with fluorescent beads. We analyzed macrophages that phagocytosed beads using flow cytometry and observed a significant reduction in the percentage of phagocytic macrophages following exposure to conditioned medium from PBMCs treated with poly(I:C) and kanamycin compared with the corresponding no-antibiotic control (**Supplementary fig. 9G**). These results demonstrate that the kanamycin-associated changes in immune responses can propagate to a downstream immune effector function and immune cell composition *in vivo*.

## Discussion

In this study, we demonstrated that antibiotics exert prevalent alterations of immune cell function. Using PBMCs and cytokine secretion as a functional, sensitive, and quantitative readout, we mapped immune activation programs across diverse cell types and in response to different ligands. We demonstrated the capacity of antibiotics to induce (*e.g.,* kanamycin, amikacin) or suppress (*e.g.,* ciprofloxacin) immune responses. Our observation that distinct antibiotics selectively modified PBMC responses to specific ligands supported the hypothesis that these effects reflect targeted rewiring of immune activation programs. A caveat of our screen is that PBMCs lack key immune compartments, such as neutrophils, which will need to be addressed in future studies. Dose–response analyses revealed that the immunomodulatory effects occurred at concentrations within or near physiologically relevant ranges^47^. Moreover, our comprehensive screen suggests class-level patterns of immunomodulation, with antibiotics from the same class eliciting similar immune effects. For example, aminoglycosides that target the bacterial ribosome may bind and impact the host mitochondrial^48,49^, all induced cytokine secretion in poly(I:C) stimulated PBMCs. Fluoroquinolones that target bacterial DNA topoisomerases and have previously been associated with perturbations of mitochondrial DNA homeostasis^50,51^, had similar impact on LPS- and poly(I:C)-stimulated PBMCs. Some antibiotics did not elicit similar immunomodulatory effects compared to compounds within the same class (*e.g.,* streptomycin compared to other aminoglycosides). Our screen included also antimalarial compounds with comparatively strong effects on cytokine production, as they directly engage pathways conserved in human immune cells, including lysosomal function, mitochondrial activity, and cellular metabolism^52,53^. These associations between classes of antibiotics and similar immunomodulatory effects provide a conceptual framework rather than establishing the mechanism responsible for the absence or presence of an effect for each individual compound. Furthermore, our screen was performed using a single donor, which raises the possibility for donor-specific effects and that compounds with immunomodulatory activity in other individuals may therefore not have been identified in our initial screen. Thus, lack of activity in this screen should not be interpreted as evidence that a compound lacks immunomodulatory activity. Nevertheless, these findings suggest antibiotics as previously underappreciated, ligand- and class-specific modulators of human immune function.

Our assays, utilizing primary human PBMCs, enabled the detection of integrated immune responses that more closely approximate *in vivo* immune coordination. The variation between donors provided a measure by which we have prioritized antibiotics based on the degree to which these effects are general or vary between individuals. To decipher the impact of antibiotics on immunometabolism we focused on ciprofloxacin that exhibited robust patterns across donors and consistent with other fluoroquinolones (*e.g.,* levofloxacin). Using scRNA-seq, we disentangled cell type-specific responses to the immunomodulatory effects of ciprofloxacin, including differential impact of ciprofloxacin on mitochondrial gene expression, most notably in T cells. These findings prompted us to use ciprofloxacin to study the mechanisms by which antibiotics may alter immunometabolism. We utilized a model of T cell expansion from PBMCs and provided several layers of evidence into the involvement of the mitochondria in ciprofloxacin mediated immunomodulation of T cells, also in concentrations which approximate the reported human plasma *C*max. Using bulk-RNA seq of T cells we detected that ciprofloxacin modified the expression levels of ETC genes encoded from the mitochondria, oxidative stress genes and proliferation. Given the prominent changes in mitochondrial gene expression, we sought to determine whether these transcriptional alterations were accompanied by a functional mitochondrial phenotype. T cell activation is associated with a shift toward aerobic glycolysis^54^. We therefore compared the effects of ciprofloxacin on T cell viability in glucose-versus galactose-containing media. Unlike glucose, galactose does not support ATP production through glycolysis, thereby increasing cellular dependence on mitochondrial oxidative phosphorylation for energy production^55^. T cells exhibited increased sensitivity to ciprofloxacin, consistent with ciprofloxacin-induced impairment of mitochondrial function. Finally, we performed Seahorse extracellular flux analysis and demonstrated that ciprofloxacin directly alters mitochondrial respiration in T cells. Together, these findings provide functional evidence that physiological-relevant concentrations of ciprofloxacin induced immunomodulation that is mediated by alterations to mitochondrial respiratory function.

Our study provides a systematic charaterization of interactions across antimicrobial compounds and immune contexts. Previous studies have established important examples of host-directed effects of ciprofloxacin and other antimicrobials. Ciprofloxacin was shown to alter mitochondrial DNA replication by inhibiting mitochondrial Top2β^56^, while AIFM1 (apoptosis induced associated factor 1) and IDH2 (isocitrate dehydrogenase 2) have also been proposed^51^. Ciprofloxacin was also shown to impact mitochondrial and immune-cell functions^57^. A recent high-resolution respirometry study in permeabilized HEK293T cells showed that ciprofloxacin directly perturbs mitochondrial respiration, increasing leak respiration and reducing respiratory capacity under both NADH- and succinate-linked conditions^58^). Aminoglycosides were shown to enhance antiviral immunity independently of the microbiota^59^, tetracyclines can perturb mitochondrial function and promote disease tolerance during infection^60^, and macrolides have been reported to modulate inflammatory pathways, including NF-κB and MAPK signaling^61^. While these studies establish that individual antibiotics can engage host pathways, our comprehensive analysis demonstrated antibiotic-, stimulus-, donor-, and cell-type-dependent effetcs, with direct alterations of immune-cell metabolism emerging as an important component of the response. These suggest convergent patterns with a shared molecular or cellular targets rather than compound-specific off-target effects.

To establish the significance of our findings in an *in vivo* setting, we performed a short stimulation of mice followed by a single dose of one of our hits, kanamycin. Our *in vivo* finding of a dose dependent increase in immune activation by kanamycin mirrored our observations in human PBMCs, indicating that these may extend also to the clinical setting. We speculate that the short time frame of immune activation precludes the involvement of the microbiome in these effects. Indeed, in a recent study, other aminoglycosides were shown to enhance host resistance to several viral pathogens, including herpes simplex virus, influenza A virus, and Zika virus, evident also in germ-free mice^28^. We further demonstrated the effects of immunomodulatory antibiotics, irrespective of their antimicrobial activity, using exposure to heat-killed bacteria and live viral infection. Our findings provide a framework for future studies that will examine the impact of immunomodulatory effects of antibiotics on bacterial clearance by the immune system during treatment. Dissecting these interactions during *in vivo* infection entails several challenges. First, separating the direct impact of antibiotics on the immune system from their antimicrobial activity. Second, pharmacokinetics of antibiotics, including cellular permeability and intracellular accumulation, are likely to influence the impact of antibiotic on immune cell function, and activity against intracellular pathogens^62^. Third, antibiotics may perturb the microbiome or cause organ-specific host toxicity, that will indirectly impact host immunity (*e.g.,* aminoglycosides can cause nephrotoxicity through renal accumulation^63,64^). These effects could be disentangled using antibiotic resistant bacteria, and infection performed *in vivo* over a short time frame, minimizing organ toxicity and microbiome disruption. Such approaches could disentangle drug–immune interactions from antimicrobial activity and determine how their interplay ultimately shapes immune-mediated pathogen clearance during antimicrobial treatment. Several studies already support this framework. For example, bedaquiline was shown to reprogram macrophages into potent bactericidal phagocytes, and enhance intracellular killing of *Mycobacterium tuberculosis* strain that is insensitive to bedaquiline^30^. Another study used *E. coli* strains carrying resistance plasmids to ribosomal-targeting antibiotics, demonstrated that tetracycline antibiotics perturb the mitochondrial electron transport chain and mediate disease tolerance to infection^26^. Finally, ciprofloxacin treatment was shown to disrupt adaptive immune responses require for protection from enteropathy in a mouse infection model of *Salmonella* Typhimurium^65^

Antibiotics have been extensively studied for their antimicrobial mechanisms, effects on the microbiome and their general toxicity in eukaryotic cells. Here, we provide comprehensive and mechanistic insight into the impact of antibiotics on mitochondrial reprogramming during immune cell activation. We suggest that antibiotic-specific molecular interactions can engage distinct host pathways to impact functional immune responses. These findings support a model for antibiotic treatment as a three-component system involving the drug, microorganisms, and host immune responses. The net consequences of antibiotic treatment may therefore reflect a complex interplay between well-established drug–microbe and microbiome–host interactions and direct antibiotic–immune effects, including the immunometabolic reprogramming demonstrated here, which may in turn influence immune function and ultimately microbial clearance. We foresee that our results will redefine the understanding of antibiotic impact on immunity, opening entirely new avenues for therapeutic intervention in infection and inflammation.

## Materials and Methods

### PBMCs extraction and handling

#### Ethics

The study was conducted in accordance with the Declaration of Helsinki and national and institutional standards and approved by the Weizmann Institute of Science Ethics Committee-approval.

#### Healthy donors

Leukocyte-enriched fractions from healthy donors were obtained from the national blood bank (MDA, Israel). Blood bank samples were fully anonymized, with no donor-identifiable information, ensuring confidentiality and compliance with ethical regulations. No other information other than blood type is provided with the samples. PBMCs were isolated, aliquoted at 20 × 10^6 cells per tube, and cryopreserved as previously described^66^. For each experiment, PBMCs were thawed (the number of donors is indicated for each experiment), counted, and assessed for viability using ViaStain™ AOPI Staining Solution (Revvity, cat. no. CS2-0106, 5 ml). Cells were resuspended in medium consisting of RPMI 1640 with L-glutamine, supplemented with 10% heat-inactivated fetal bovine serum and 1 mM sodium pyruvate (from now on RPMI+). PBMCs were plated at 1 × 10^5 cells per well in U-bottom, untreated 96-well plates (Greiner Bio-One, cat. no. 650185) and incubated at 37 °C with 5% CO2 for 1.5 h prior to further treatment.

#### Patient donors

Peripheral blood (10 mL) was collected from 20 patients admitted to Ichilov Medical Center (Tel Aviv, Israel) who presented with symptoms of infectious disease. All participants provided written informed consent prior to inclusion, and the study was approved by the institutional ethics committee. Donor-identifiable information, including names and identification numbers, was retained exclusively by the clinical center and was not accessible to the investigators. Patient metadata are provided in Table 3. Whole blood samples were diluted 1:1 with phosphate-buffered saline lacking calcium and magnesium (from now on PBS -/-) supplemented with 2% fetal bovine serum and gently mixed. Samples were centrifuged at 800 × g for 10 min at room temperature (15–25 °C) with the brake disengaged. Following centrifugation, the buffy coat fraction (approximately 5 mL) was carefully transferred to a new 50 mL conical tube, diluted with 20 mL PBS -/-, and layered over Lymphoprep (Alere Technologies AS) at a diluted blood–to– Lymphoprep ratio of 1:0.54. Gradients were centrifuged at 630 × g for 30 min at room temperature using low acceleration (setting 1) and minimal braking (setting 1). After centrifugation, the peripheral blood mononuclear cell (PBMC) layer at the plasma-Lymphoprep interface was collected into a new 15 mL tube. Cells were washed in PBS-/-containing 1% FBS and 2.5 mM EDTA, followed by centrifugation at 630 × g for 15 min at 4 °C. The supernatant was discarded, and the cell pellet was resuspended in 500 µL RPMI+ medium. Cell concentration and viability were determined using AOPI viability staining. Patient-derived PBMCs were subsequently cultured and treated under the same conditions as healthy donor PBMCs.

### Human PBMCs screen for the immunomodulatory effect of drugs

Each condition in the screen (naïve, LPS or poly(I:C) was performed using PBMCs derived from a single donor. Cryopreserved PBMCs were thawed, counted, and resuspended in RPMI+ medium. Cells were plated at a density of 1 × 10^5 cells per well in untreated U-bottom 96-well plates and incubated at 37 °C with 5% CO₂ for 1.5 h prior to stimulation. Following preincubation, cells were stimulated with lipopolysaccharide (LPS; 50 pg/mL; Sigma-Aldrich, cat. no. L3024-5MG) or polyinosinic–polycytidylic acid (poly(I:C); 50 µg/mL; InvivoGen, cat. no. tlrl-pic). For Supplementary Fig. 1a,b-see stimulant concentration in the figure. Cells were gently mixed and incubated at 37 °C with 5% CO₂ for an additional 1 h. Subsequently, test drugs were added at the concentrations indicated in Supplementary Table 2, followed by gentle mixing and incubation for either 4 h or 24 h at 37 °C with 5% CO₂. At the end of each incubation period, plates were centrifuged at 500 × g for 5 min. Supernatants were carefully collected without disturbing the cell pellets and stored at -80 °C until analysis of cytokine and signaling molecule activity using HEK-Blue reporter cell assays (see HEK-Blue reporter cell at Supplementary Table 1).

### Signaling molecules and cytokines measurement by HEK-Blue reporter cells

HEK-Blue reporter cell lines (listed in Supplementary Table 1) were cultured and maintained according to the manufacturer’s instructions (InvivoGen). To assess cytokine production, 20 µL of culture supernatant (from a total of 180 µL) from PBMCs or splenocytes, was added to each HEK-Blue reporter cell line. Reporter activity was quantified according to the manufacturer’s protocol (InvivoGen) using Quanti-Blue detection solution (InvivoGen, cat. no. rep-qbs). Absorbance was measured at OD630 nm using FLUOstar Omega microplate reader (BMG Labtech).

### RNA-seq experiment and analysis

PBMCs from a single donor were thawed, stimulated, and treated with antibiotics as described in the ‘Human PBMC screen for the immunomodulatory effect of drugs’ method section. See the used drug concentration in each experiment. For PBMCs bulk RNAseq, samples were collected at the following time points after treatment: 0.5, 1, 2, 4, 6, and 24 h. Naïve (unstimulated) samples without antibiotic treatment were collected only at 0.5 h and 24 h. At each indicated time point, cells were centrifuged (500 × g, 22 °C, 5 min), the supernatant was discarded, and pellets were washed with PBS -/-. Cells were then resuspended in PBS -/-containing 5 mM EDTA and incubated for 5 min at room temperature, followed by gentle pipetting to detach adherent cells. Samples were centrifuged again (500 × g, 22 °C, 5 min) and resuspended in 350 µL RLT lysis buffer (RNeasy Mini Kit, QIAGEN, cat. no. 74104) supplemented with β-mercaptoethanol (10 µL per 1 mL RLT buffer). Lysates were stored at -80 °C until RNA extraction.

For bulk RNA-seq, T cells from one donor were activated and expanded (see method section ‘T cell activation and expansion’), then treated with ciprofloxacin at the indicated concentrations for 4 days. The cells were then centrifuged (500 × g, 4 °C, 5 min), the supernatant was discarded, and the pellets were washed with PBS -/-. Cells were then resuspended in 350 µL RLT lysis buffer (RNeasy Mini Kit, QIAGEN, cat. no. 74104) supplemented with β-mercaptoethanol (10 µL per 1 mL RLT buffer). Lysates were stored at -80 °C until RNA extraction.

For mice splenocyte RNAseq, spleens were harvested from individual mice and mechanically dissociated through a 70 mm cell strainer into 50 ml conical tubes using a sterile syringe plunger. The strainer was washed with 3 ml of ice-cold PBS +/+ supplemented with 2% fetal bovine serum (FBS). Cell suspensions were centrifuged at 500 g for 5 min at 4 °C. Pelleted single-cell suspensions were resuspended in 2 mL of cryopreservation medium (RPMI-1640 supplemented with 30% FBS and 10% DMSO). The 2 mL suspension from each mouse spleen was distributed equally into four cryovials (0.5 ml per vial) and kept in −150 °C freezer for long-term storage. Upon thawing cells were washed with RPMI+ media, then resuspended in 350 µL RLT lysis buffer (RNeasy Mini Kit, QIAGEN, cat. no. 74104) supplemented with β-mercaptoethanol (10 µL per 1 mL RLT buffer). Lysates were stored at -80 °C until RNA extraction.

Total RNA was extracted using the RNeasy Mini Kit (QIAGEN, cat. no. 74104), including on-column DNase digestion (QIAGEN, cat. no. 79254), according to the manufacturer’s instructions. RNA concentration was determined using the iQuant™ RNA HS Assay Kit (ABP Biosciences, cat. no. N017), and RNA integrity was assessed using the TapeStation system with High Sensitivity RNA ScreenTape (Agilent, cat. no. 5067-5579) and High Sensitivity RNA ScreenTape Sample Buffer (Agilent, cat. no. 5067-5580). RNA samples were stored at −80 °C until further processing.

RNA-seq libraries were prepared according to the CEL-Seq2 protocol^67^, with a minor modification-purified bulk RNA was used as input (up to 1 ng RNA per library). Libraries were sequenced on an Illumina NovaSeq X instrument using the NovaSeq X Series 1.5B Reagent Kit (100 cycles; Illumina, cat. no. 20104703), PBMC libraries had a total of approximately 1250 million reads, with a median depth of approximately 8 million reads per library. T cell libraries had a total of approximately 302 million reads, with a median depth of approximately 11 million reads per library. Mice splenocyte libraries had a total of approximately 267 million reads, with a median depth of approximately 11 million reads per library.

The cel-seq pipeline (https://github.com/yanailab/CEL-Seq-pipeline) was used for sample demultiplexing, genome alignment (hg19 for PBMCs and T cells, mm10 for mouse experiments), and gene counting to generate an expression matrix for each dataset. One sample (*LPS_CIP_4h_rep3*) was excluded from the PBMCs analysis (< 500k reads), and three samples were excluded from the T cells analysis due to low coverage (<200K reads; replicates 2,4, and 5 of the control condition without ciprofloxacin).

For each dataset, the expression matrix was normalized to a library-size factor by dividing the total number of reads for each sample by the median across all samples in that experiment. Normalized data was transformed to a log2 scale, and a minimal expression threshold of 2 was set for downstream. Replicate samples of each condition were averaged (3-7 replicates per condition). Preceding PCA analysis, genes were centered and normalized to a mean of 0 and a standard deviation of 1. For the PBMCs dataset, to identify genes up- or down-regulated by antibiotics, we calculated the difference between the integrals of each gene in the absence of antibiotics and those under each antibiotic treatment over time. The differences across all genes were approximately normally distributed, with a median of 0 and a standard deviation of 6.6-10.7 (depending on the condition). Genes with more than 3 standard deviations above the mean were defined as up- or down-regulated due to antibiotic treatment. KEGG pathway enrichment analysis of significantly differentially expressed genes was performed using a hypergeometric test with FDR correction for multiple testing (10%FDR). For the mouse dataset, DESeq^68^ was used to identify differentially expressed genes (10%FDR) between poly(I:C) only and poly(I:C) with kanamycin treatment at different concentrations.

### Single-cell RNA libraries preparation

PBMCs from a single donor were thawed, stimulated, and treated with antibiotics as described in the ‘Human PBMC screen for the immunomodulatory effect of drugs’ method section. Ciprofloxacin concentrations are indicated in the text and in Fig.4. After 24h, the cells were centrifuged (500 × g, 22 °C, 5 min), the supernatant was discarded, and the cells were washed with PBS -/-. Cells were then resuspended in PBS -/- containing 5 mM EDTA and incubated for 5 min at room temperature, followed by gentle pipetting to detach adherent cells. Samples were centrifuged again (500 × g, 22 °C, 5 min) and resuspended in 100ul of PBS -/- + 0.04% BSA. The cells were counted and libraries were generated using the GEM-X Universal 3′ Gene Expression v4 4-plex assay with on-chip multiplexing (OCM; 10x Genomics), according to the manufacturer’s instructions (see User Guide | CG000768 | Rev B https://www.10xgenomics.com/support/universal-three-prime-gene-expression/documentation/steps/library-prep/gem-x-universal-3-prime-gene-expression-v-4-4-plex-reagent-kits)

Libraries were sequenced on an Illumina NovaSeqX instrument using the following read configuration: Read1 28bp, Read2 90bp, Index1 10bp, Index2 10bp. yielding a total of 2448M reads across all libraries.

### Single-cell RNA-seq analysis

#### Processing

Raw sequencing data were processed using Cell Ranger (version 10.0.0; 10x Genomics). Samples within each 4-plex reaction were computationally demultiplexed based on their 10x barcode assignments. Sequencing reads were aligned to the human reference genome (GRCh38), and cell-associated barcodes and unique molecular identifiers (UMIs) were identified and quantified using the Cell Ranger pipeline (cellranger multi). The resulting gene-by-cell UMI count matrices for each sample were used for downstream quality control and single-cell transcriptomic analyses. Overall, we sequenced 43,499 cells (mean 3,625 cells per sample, standard deviation 717), with a mean of 10,255 UMIs and 3,060 genes per cell, and a total of 23,335 genes detected.

#### QC filtration and normalization

Only genes with at least one UMI count detected in at least one cell were used. Cells with a high percentage of mitochondrial genes (more than 20%, 43 cells) were excluded. All cells had a total count of at least 500 UMIs. Data was normalized to a library size factor, with factors calculated by dividing the total UMI counts in each cell by the median of the total UMI counts across all cells. Normalized data was log2-transformed. Cell cycle and ribosomal genes were filtered out prior to the selection of variable genes. Variable genes were selected based on fitting the data to a simple noise model based on the gene’s mean expression and dispersion (coefficient of variance) across all single cells.

#### Clustering and data analysis

Principal component analysis (PCA) was performed on the variable genes, and the first 34 PCs were selected for downstream analysis based on the inflection point of the curve of the variance explained by each successive principal component. tSNE in the PC space was calculated for the visualization of the data. The Euclidian distance in the PC space and the Jaccard similarity were used to build a k-nearest neighbor (KNN)-graph with k=30. Unsupervised clustering of the single cells was done using Louvain community detection. We used an iterative implementation of the Louvain method^69^ to optimize modularity-like quality function by iterating the Louvain algorithm until convergence. Cluster identity was inferred using cluster-specific genes and cell-type marker genes. Next, we re-cluster each cell type to the subset level using Louvain algorithm. Overall, we obtained 46 immune subsets and 2 non-immune subsets (Megakaryocytes/Platelets and Erythroid; **Supplementary fig. S5A-B**).

### Mice experiments

Female C57BL/6 mice (7–9 weeks old) were purchased from Envigo and housed in the pathogen-free animal facility of the Weizmann Institute of Science under standard specific pathogen-free (SPF) conditions, with ad libitum access to food and water. All animal experiments were conducted in accordance with institutional guidelines and were approved by the institutional IACUC committee. For *in vivo* stimulation, mice were injected intravenously with PBS (control) or poly(I:C) (150 µg per mouse), followed by subcutaneous injection of PBS (control) or kanamycin at doses of 500, 700, or 1,100 mg/kg (indicated in each figure). Four hours after injection, blood was collected via retro-orbital bleeding into EDTA-coated tubes. Blood samples were centrifuged at 2,000 × g for 10–15 min at 4 °C, and plasma (upper phase) was collected and stored at -80 °C until analysis.

For splenocyte isolation, spleens were harvested from naïve mice and mechanically dissociated by gentle crushing through a 70 µm cell strainer into 5 mL of PBS + 2% FBS per spleen. Cell suspensions were centrifuged at 500 × g for 5 min at 4 °C. Pellets were resuspended in 2 mL of 1× freezing medium consisting of RPMI 1640 supplemented with 30% FBS and 10% dimethyl sulfoxide (DMSO), aliquoted, and cryopreserved. For experiments, splenocytes were thawed, counted, and assessed for viability using AO/PI stain, resuspended in RPMI+ medium, and plated at a density of 1 × 10^5 cells per well in untreated U-bottom 96-well plates. Cells were incubated at 37 °C with 5% CO₂ for 1.5 h prior to stimulation. Following preincubation, cells were stimulated with LPS (1 µg/mL) or poly(I:C) (50 µg/mL) and incubated for 1 h at 37 °C with 5% CO₂. Subsequently, drugs were added at the concentrations indicated in Supplementary Fig.9. The cells were incubated for 24 h at 37 °C with 5% CO₂. Plates were then centrifuged at 500 × g for 5 min. Supernatants were carefully collected without disturbing the cell pellets and stored at -80 °C until analysis of cytokine and signaling molecule activity using HEK-Blue reporter cell assays specific for mouse cytokines (Supplementary Table 1), as described in the ‘Signaling molecules and cytokines measurement by HEK-Blue reporter cells*’* section.

### Heat-inactivation *E. coli* treatment

PBMCs from a healthy donor were thawed, counted, and assessed for viability using AO/PI stain, resuspended in RPMI+ medium, plated at a density of 1 × 10^5 cells per well in untreated U-bottom 96-well plates, and rested for 1.5h at 37 °C with 5% CO₂. An overnight culture of *Escherichia coli* (K2 strain) was quantified by measuring optical density at 600 nm (OD600). To validate the multiplicity of infection (MOI), bacterial colony-forming units (CFU) were determined by plating serial dilutions on LB agar plates overnight at 37 °C prior to inactivation. Bacteria were heat-killed by incubation at 65 °C for 60 min; complete loss of bacterial viability was confirmed by plating post-heat-inactivation bacteria on LB agar overnight. PBMCs were left unstimulated or challenged with heat-killed E. coli at an MOI of 0.5, followed by the addition of ciprofloxacin at indicated concentrations (0, 13.5, 52.5, 105, or 210 μm) for 24 h. PBMCs were then centrifuged (500g for 5min), and supernatants were collected and stored at −80 °C. Changes in cytokine levels were detected as described in the “Signaling molecules and cytokine measurement by HEK-Blue reporter cells” methods section.

### Viral infection

PBMCs from three donors were thawed, counted, and assessed for viability using AO/PI stain, resuspended in RPMI+ medium, plated at a density of 1 × 10^5 cells per well in untreated U-bottom 96-well plates, and incubated overnight at 37 °C with 5% CO₂. Cells were subsequently infected with the human cytomegalovirus (HCMV) TB40 strain at a multiplicity of infection (MOI) of 10. This recombinant strain expresses mNeonGreen linked to immediate-early genes (UL122/UL123), mTagBFP2 fused to early genes (UL112/UL113), and mCherry fused to a late viral gene (UL48A), enabling temporal resolution of viral gene expression^45^. PBMCs in the presence of virus were centrifuged at 800 × g for 1 h at room temperature, followed by three washes with RPMI+ medium. Antibiotics were then added at the indicated concentrations. Cells were gently mixed by pipetting and incubated for 18 h at 37 °C with 5% CO₂. Plates were then centrifuged at 500 × g for 5 min. Supernatants were carefully collected without disturbing the cell pellets and stored at -80 °C until analysis of cytokine and signaling molecule activity using HEK-Blue reporter cell assays (see HEK-Blue reporter cell at Supplementary Table 1). For validation of viral infection, cells were centrifuged (500 × g, 22 °C, 5 min), the supernatant was discarded, and pellets were washed with PBS -/-. Cells were resuspended in PBS -/- containing 5 mM EDTA and incubated for 5 min at room temperature, followed by gentle pipetting to detach adherent cells. Samples were analyzed directly by flow cytometry using a CytoFLEX instrument (Beckman Coulter). Data analysis was performed using FlowJo v10.7.1.

### Phagocytosis assays

PBMCs from a single donor were thawed, stimulated, and treated with kanamycin (40 μg/ml) as described in the “Human PBMC screen for the immunomodulatory effect of drugs” methods section. After 24 h, cell were centrifuged (500 × g, 22 °C, 5 min), and the resulting supernatant (conditioned medium) was collected and stored at -80 °C until further use.

THP-1 cells were seeded into tissue culture-treated 96-well flat-bottom plates (1*10^5 cells/well) in RPMI+ medium and differentiated into macrophage-like cells using 100 ng/ml phorbol 12-myristate 13-acetate (PMA) for 48 h. The culture medium was then replaced with PMA-free RPMI+ medium for a 48 h resting period. Following resting, the medium was replaced with thawed conditioned medium and incubated for 24 h. Fluorescent latex beads (Sigma-Aldrich, Cat# L1030) were added at a bead-to-cell ratio of 200:1, followed by centrifugation (500 × g, 22 °C, 5 min). Cells were incubated with beads for 3 h at 37 °C, washed with PBS-/-and resuspended in PBS-/- containing 5 mM EDTA for 5 min at room temperature. Adherent cells were detached by gentle pipetting, and samples were acquired directly on a FACSAria III flow cytometer (BD Biosciences) using BD FACSDiva software. Data analysis was performed using FlowJo software (v10.7.1, BD Biosciences).

### Enzyme-linked immunosorbent assay (ELISA)

PBMCs culture supernatants were harvested following centrifugation (500 × g, 22 °C, 5 min) and stored at -80 °C until analysis. Secreted levels of TNF-α (BioLegend, 430201), IFN-γ (BioLegend, 430101), IFN-β (BioLegend, 449504), and IL-6 (BioLegend, 430501) were quantified using commercial sandwich ELISA kits according to the manufacturers’ protocols. Mouse plasma cytokines were measured using ELISA kits specific for mouse IFN-β (BioLegend, 439404) and IFN-γ (BioLegend, 430804), according to the manufacturer’s instructions.

### T cell activation and expansion

A tissue culture-treated 12-well plate was coated with anti-CD3 (5 µg/µl, BD Biosciences BD567118) in PBS-/- for 2-4 h at 37°C with 5% CO_2_ incubator. Following incubation, the coating solution was removed and PBMCs were seeded at a density of 2*10^6^ cells/ml in T cell medium containing RPMI 1640 with HEPES (Thermo Fisher Scientific, 52400-025) supplemented with 10% FBS,1mM sodium pyruvate,1% penicillin/streptomycin, 2mM L-glutamine, 1% non-essential amino acid solution, 0.05mM β-mercaptoethanol, and recombinant human IL-2 (250 IU/ml, Biotest 202-IL-010). Anti-CD28 (5 µg/µl, BD Biosciences 555725B, was added before incubation for 96 h.

After 96 h, cells were supplemented with fresh T cell medium at a 1:1 ratio and transferred to a tissue culture-treated 6-well plate. Every two days thereafter, cells were supplemented again with fresh medium at a 1:1 ratio and transferred to new 6-well plates or 10 cm tissue culture dishes.

### Flow cytometry for T cells

Samples were washed with PBS-/- and incubated with Fc block (BD Biosciences 564220,) diluted in PBS-/- to prevent nonspecific antibody binding. For activation assessment, T cells were stained with CD134 (Biolegend 563663, dilution 1:100) and CD137 (Biolegend 309830 dilution 1:100). Cells were then washed and stained with Fixable Viability Stain 450 (BD Biosciences, FVS450) for the discrimination of live/dead cells. Surface antigens were stained using CD3-FITC (UCHT1 clone, BioLegend 300406) at dilution of 1:100 dilution, CD4 (SK3 clone BD Biosciences, BD566915) at dilution of 1:100, CD8 (SK1 clone, BD Biosciences BD565310) at dilution of 1:100 dilution. Cells were subsequently fixed and permeabilized using the Cytofix/Cytoperm™ kit (BD Biosciences). For intracellular staining, Ki-67-PE (B56 clone, BD Biosciences BD567719), at 1:20 dilution. DRAQ5 (BioLegend, 424101, final con. 15mM) was added for DNA content staining. Absolute cell numbers were determined using Precision Count Beads (BioLegend, 424902) according to the manufacturer’s instructions.

Flow cytometry data were acquired on a FACSAria™ III flow cytometer (BD Biosciences) using BD FACSDiva™ software. Data analysis and image generation were performed using FlowJo v10.7.1.

### Glucose-Galactose assay

T cells derived from PBMCs were cultured in T cell medium containing 11mM glucose for 7 days. On day 7, cells were treated with ciprofloxacin (13.5μM or 105 μM) and incubated for an additional 3 days in the same medium. On day 10, cells were counted and adjusted to equal cell concentration, centrifuged, and washed with PBS +/+. An equal number of cells from each condition were then cultured for 24 h in T cell medium containing either 11mM glucose (Thermo Fisher Scientific 11879020,) or galactose (Sigma PHR1206). As a positive control for mitochondrial inhibition, cells were treated for 24 h with 50nM rotenone, antimycin A, or oligomycin.

#### Seahorse mitochondrial stress assay

T cells derived from 3 PBMCs donors were cultured in T cell medium (see ‘T cell activation and expansion’ method section) with one modification-glucose concentration was 5.5 mM for 7 days. On day 7, cells were treated with ciprofloxacin (13.5μM 50 μM or 105 μM) and incubated for an additional 3 days in the same medium. On day 11, cells were counted and adjusted to equal cell concentration, centrifuged, and washed with PBS -/-. An equal number of 1.5 × 10^5^ live cells per well were seeded onto poly-D-lysine coated Seahorse plates, with 3-6 technical replicate wells per condition in Seahorse XF RPMI medium supplemented with glucose, pyruvate, and L-glutamine according to the manufacturer’s instructions. Oxygen consumption rate (OCR) and extracellular acidification rate (ECAR) were measured using a modified Mito Stress Test on the Seahorse XF Pro Analyzer. Specifically, mitochondrial respiration parameters were assessed following sequential injection of oligomycin A, carbonyl cyanide-p-trifluoromethoxyphenylhydrazone (FCCP), and rotenone/antimycin A. Basal OCR and ECAR were calculated from measurements obtained before oligomycin A injection. Proton leak was determined after injection of oligomycin A, an ATP synthase/complex V inhibitor. Maximal respiration was calculated as the highest OCR measurement obtained following two sequential FCCP injections to ensure maximal respiratory capacity. Non-mitochondrial oxygen consumption was determined after injection of rotenone, a complex I inhibitor, and antimycin A, a complex III inhibitor. Final compound concentrations were as follows: oligomycin A, 1.5 μM; FCCP, 1 μM per injection; rotenone, 0.5 μM; and antimycin A, 0.5 μM. The OCR/ECAR ratio was calculated for each well and averaged across technical replicate wells for each condition.

### qPCR mtDNA copy number

T cells from PBMCs of 3 donors were exposed to either DDW or CIP at concentrations of 13.5/50/105μM for 4 days. Cells were harvested collected by centrifugation, and the resulting cell pellets were stored at - 80c until further processing. Cell pellets were thawed, and total DNA was isolated using the QIAGEN DNeasy Blood& Tissue Kit QIAGEN, 69504) according to the manufacturer’s instructions. DNA concentration was measured using NanoDrop spectrophotometer, and samples were diluted in DDW to a final concentration of 12ng/μl. The relative abundance of mitochondrial (ND2 primers; Forward: TGTTGGTTATACCCTTCCCGTACTA, Reverse: CCTGCAAAGATGGTAGAGTAGATGA) to nuclear DNA (B2M primers; Forward: GCTGGGTAGCTCTAAACAATGTATTCA, Reverse: CCATGTACTAACAAATGTCTAAAATGGT) was determined by real-time PCR using a QuantStudio™ Real-Time PCR with 384-well block (Thermo Fisher Scientific) and was calculated using the ΔCt method. Data analysis was performed using QuantStudio Design & Analysis Software 2.8.0.

## Data Availabiliy

RNA-seq data have been deposited in NCBI’s Gene Expression Omnibus (GEO) under the superseries GSE344856.

## Acknowledgments

R.A. is supported by the Israel Science Foundation (ISF grant No. 1289/22), the Joint DFG-ISF Research Grant Program (ISF grant No. 1297/25), the ISF MAVRI program (grant No. 1209/26), the Garvan Institute of Medical Research (Garvan) - WIS Collaborative Science Program, the Knell Family Center for Microbiology, the Pasteur-Weizmann challenge, and Minerva Foundation with funding from the Federal Ministry for Education and Research.

## Contributions

S.H.A and R.A conceived this work and planned the experiments. S.H.A, R.A, I.A performed experiments. S.K.L and T.Y performed Seahorse experiments and analysis. N.B.M analyzed the data. A.Z and R.B.A collected patients’ samples.

