## Supplementary_figures for "Antibiotics drive immunometabolic rewiring of human immune responses"

### Supplementary Figure 1

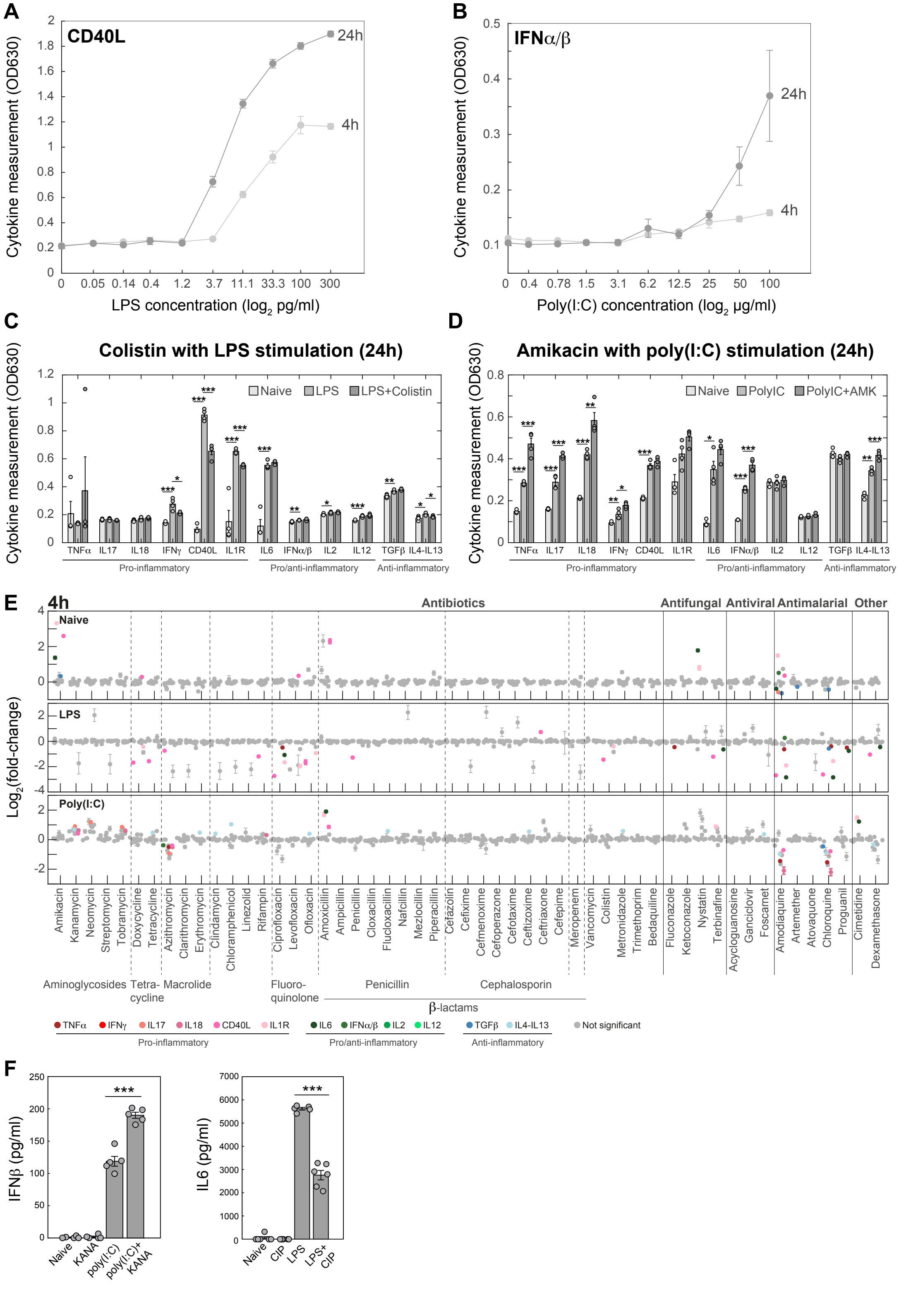

**Supplementary Figure 1: A comprehensive screen identified immunomodulatory effects of antimicrobial drugs.** (**A+B**) PBMCs were stimulated with the indicated concentrations of LPS (X-axis; A) or poly(I:C) (B) and tested for secretion of cytokines (Y-axis). (**C+D**) Bar plots indicating the effects of selected hit drugs. (**C**) PBMCs were either unstimulated (light grey bars) or stimulated with LPS (grey bars) or with LPS in the presence of colistin (dark bars) and levels of 12 cytokines (shown on the x-axis) were measured. (**D**) PBMCs were either unstimulated (light grey bars) or stimulated with poly(I:C) (grey bars) or with poly(I:C) in the presence of amikacin (dark bars) and levels of 12 cytokines (shown on the x-axis) were measured. (**E**) PBMCs from a healthy donor were either unstimulated (upper) or stimulated with LPS (middle) or poly(I:C) (lower), exposed to the indicated drugs (X-axis; drug concentrations are in Supplementary Table 2) for 4h, and tested for secretion of a panel of 12 cytokines (indicated in the colorbar). For each condition, log2(fold change) between with and without drug was calculated (Y-axis). Non-significant changes are shown in grey, and significant changes with 1%FDR fold are colored (2-sample t-test). Results are from four replicates presented as mean ± standard error. (**F**) ELISA measurements from supernatant of PBMCs treated for 24h with poly(I:C) and kanamycin (40 μg/ml) (IFNβ; left panel) or treated with LPS and ciprofloxacin (210 μM) (IL6; right panel). *P* values indicated are from a 2-sample t-test, * $p<0.05$, ** $p<0.01$, *** $p<0.001$.

#### Supplementary Figure 2

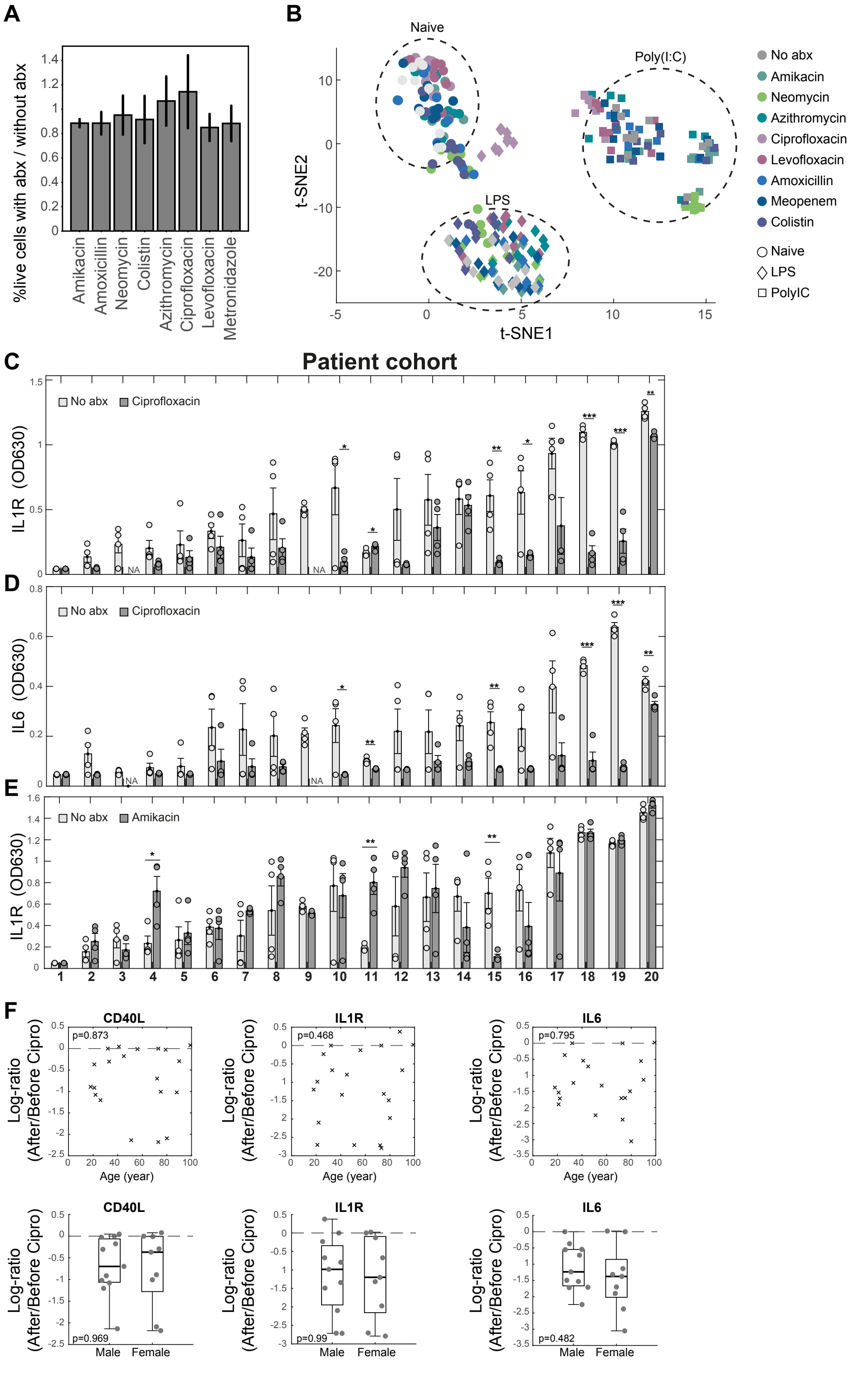


**Supplementary Figure 2: Analyzing inter-individual variation of the effects of immunomodulatory antibiotics.** (**A**) PBMCs were exposed to the indicated antibiotics for 24h (X-axis, see concentrations in

Supplementary table 2), and viability was measured and compared to no drug control treatment (Y-axis). (**B**) t-SNE representation of PBMCs from 10 individuals, treated for 24h in 3 conditions: naïve, LPS, and poly(I:C) with eight antibiotics and no treatment control (indicated in the colorbar), and measured for cytokine secretion of TNFα, IFNγ, CD40L, IL1R, IL6, and IFNα/β. (**C-E**) PBMCs from 20 patients (X-axis) were collected and exposed to ciprofloxacin (**C-D**) or amikacin (**E**) for 24h, and analyzed for IL1R (IL1α/β; **C**) or IL6 levels (**D**) for ciprofloxacin or IL1R levels for amikacin (Y-axis). (**F**) Log-ratios of cytokine levels (from patients' data in C-E), before and after ciprofloxacin treatment (Y-axis) against patient age (upper panels; X-axis) or sex (lower panels; X-axis). Associations with age and sex were assessed using linear regression models. *P* values correspond to the age or sex coefficient in the linear regression model. *P* values indicated are from a 2-sample t-test, * $p<0.05$, ** $p<0.01$, *** $p<0.001$. abx- antibiotics.

#### Supplementary Figure 3

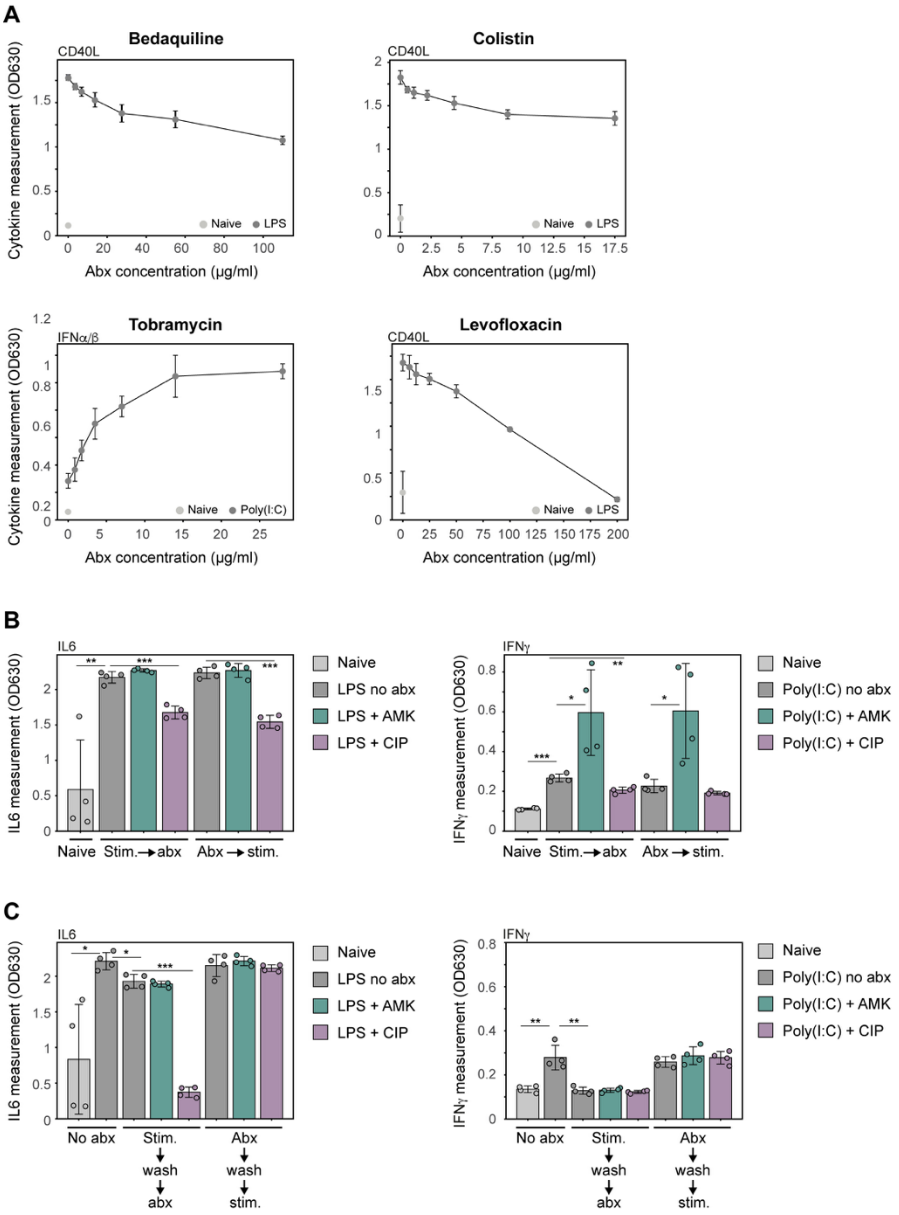


**Supplementary Figure 3: Dose-dependent and activation state-dependent immunomodulatory effects of antibiotics.** (**A**) Indicated are the cytokine measurements of PBMCs, either naïve (light grey) or stimulated (dark grey), and treated with different doses of the indicated antibiotics (X-axis). (**B**) PBMCs from a healthy donor were either stimulated with LPS (left) or poly(I:C) (right) before or after antibiotic exposure (indicated on X-axis) and measured for cytokine secretion (Y-axis). Naïve PBMCs are indicated in light grey, and stimulated PBMCs without antibiotics are indicated in dark grey. (**C**) PBMCs from a healthy donor were stimulated with LPS (left) or poly(I:C) (right), the stimulant was washed after 1h, and the indicated antibiotics were added (X-axis) and measured for cytokine secretion (Y-axis). Alternatively, PBMCs were treated with the indicated antibiotics for 1h, the antibiotics were washed, and the stimulant was added. Naïve PBMCs are indicated in light grey, and stimulated PBMCs without antibiotics are indicated in dark grey. P values indicated are from a 2-sample t-test, * $p<0.05$, ** $p<0.01$, *** $p<0.001$. AMK- amikacin, CIP- ciprofloxacin, stim- stimulant, Abx- antibiotics. Antibiotic concentrations in B-C: amikacin- 70 μg/ml, ciprofloxacin- 70 μg/ml.

### Supplementary Figure 4

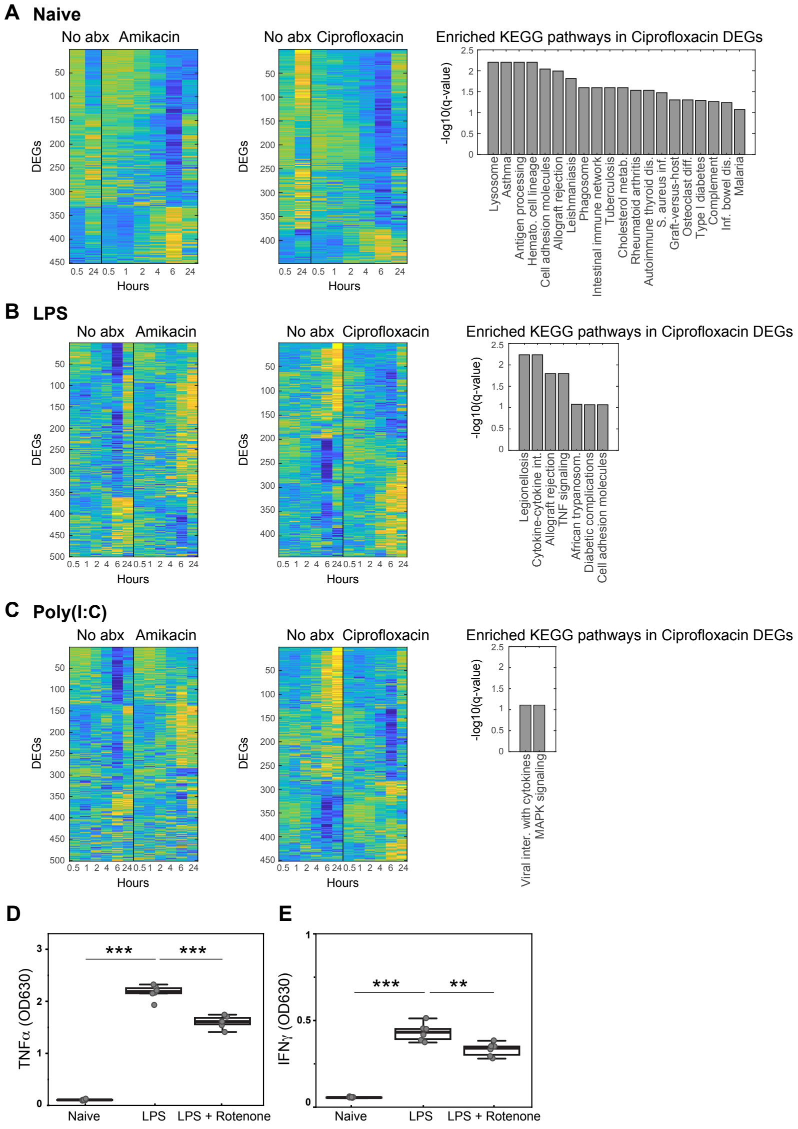


**Supplementary Figure 4: Transcriptional analysis reveals gene expression alterations by amikacin and ciprofloxacin.** (**A-C**) Heatmaps showing differentially expressed genes (DEGs) of PBMCs from a healthy donor, untreated or treated with ciprofloxacin (70 µg/ml), or amikacin (70 µg/ml) at different time points, either naïve (**A**) or stimulated with LPS (**B**) or poly(I:C) (**C**). Enriched signaling pathways are shown for ciprofloxacin-treated samples. Abx- antibiotics. (**D-E**) PBMCs from a healthy donor were kept unstimulated or exposed LPS or LPS + 50nM Rotenone (mitochondrial complex I inhibitor). Supernatants were than collected and analyzed for TNFα (**D**) or IFNγ (**E**). Box plots represent mean ± SEM. *P* values indicated are from 2-sample t-test, relative to the untreated control (naive): ** P< 0.01, ***P<0.001.

### Supplementary Figure 5

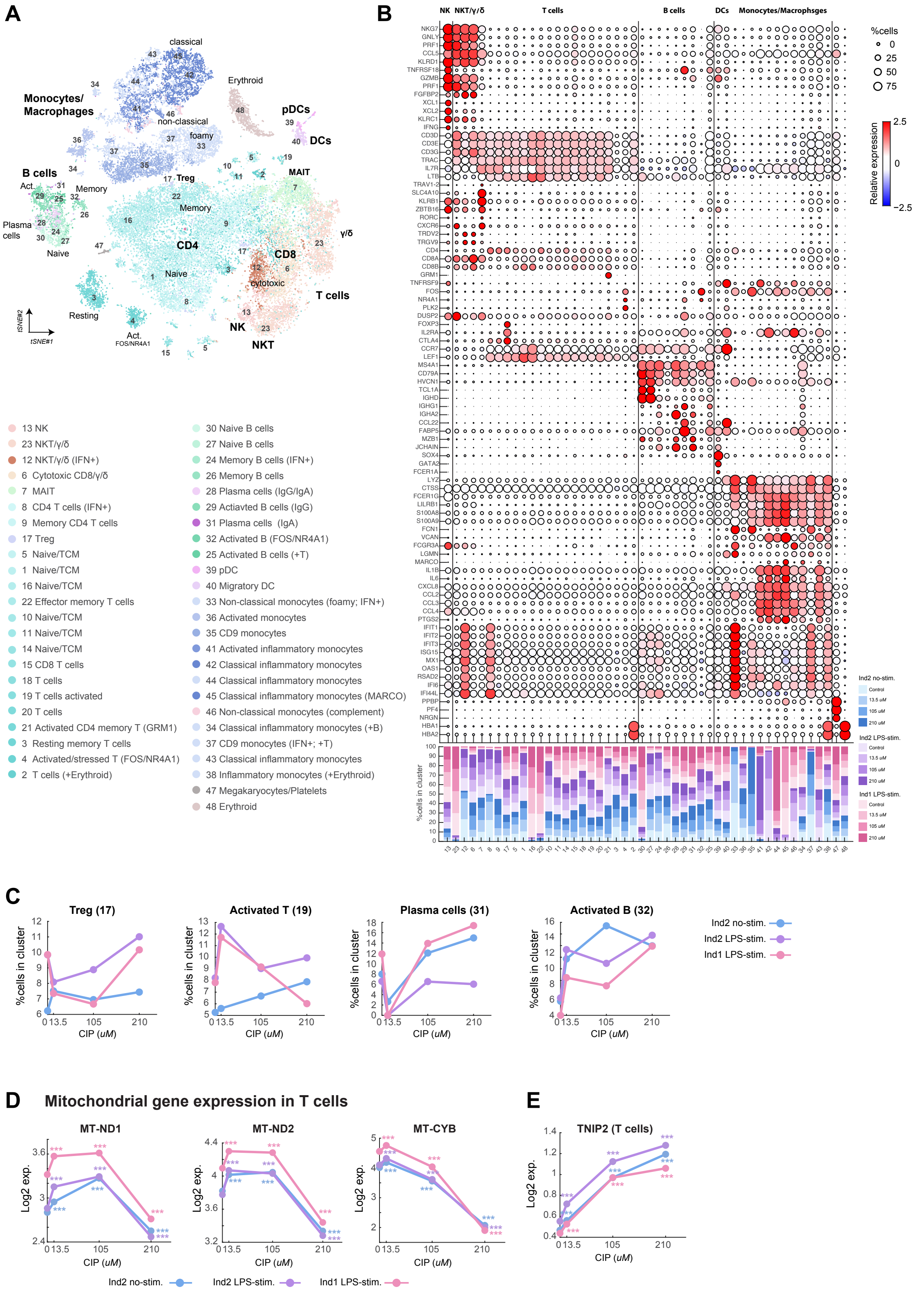

**Supplementary Figure 5: Single-cell RNA-seq reveals transcriptional changes in mitochondrial genes induced by ciprofloxacin treatment. (A)** tSNE visualization of scRNA-seq data (from Fig. 4), combining the individuals and treatments together, colored by the cell subset. Legend with numbers and cell subsets annotation is provided. **(B)** Upper panel: Dot plot showing marker gene expression levels for each cell subset. Dot size represents the percentage of cells expressing a given gene within each subset, and dot color indicates mean relative expression levels (see legend on the right). Bottom panel: Proportions of each cell subset across all conditions. **(C)** Dynamics of the percentage of cells across different ciprofloxacin concentrations per individual and stimulation within representative cell subsets. Subsets' names and numbers correspond to A. **(D-E)** Dynamics in T cells gene expression of representative mitochondrial encoded genes (**D**) or nuclear encoded gene (**E**) over different ciprofloxacin concentrations per individual and stimulation.

### Supplementary Figure 6

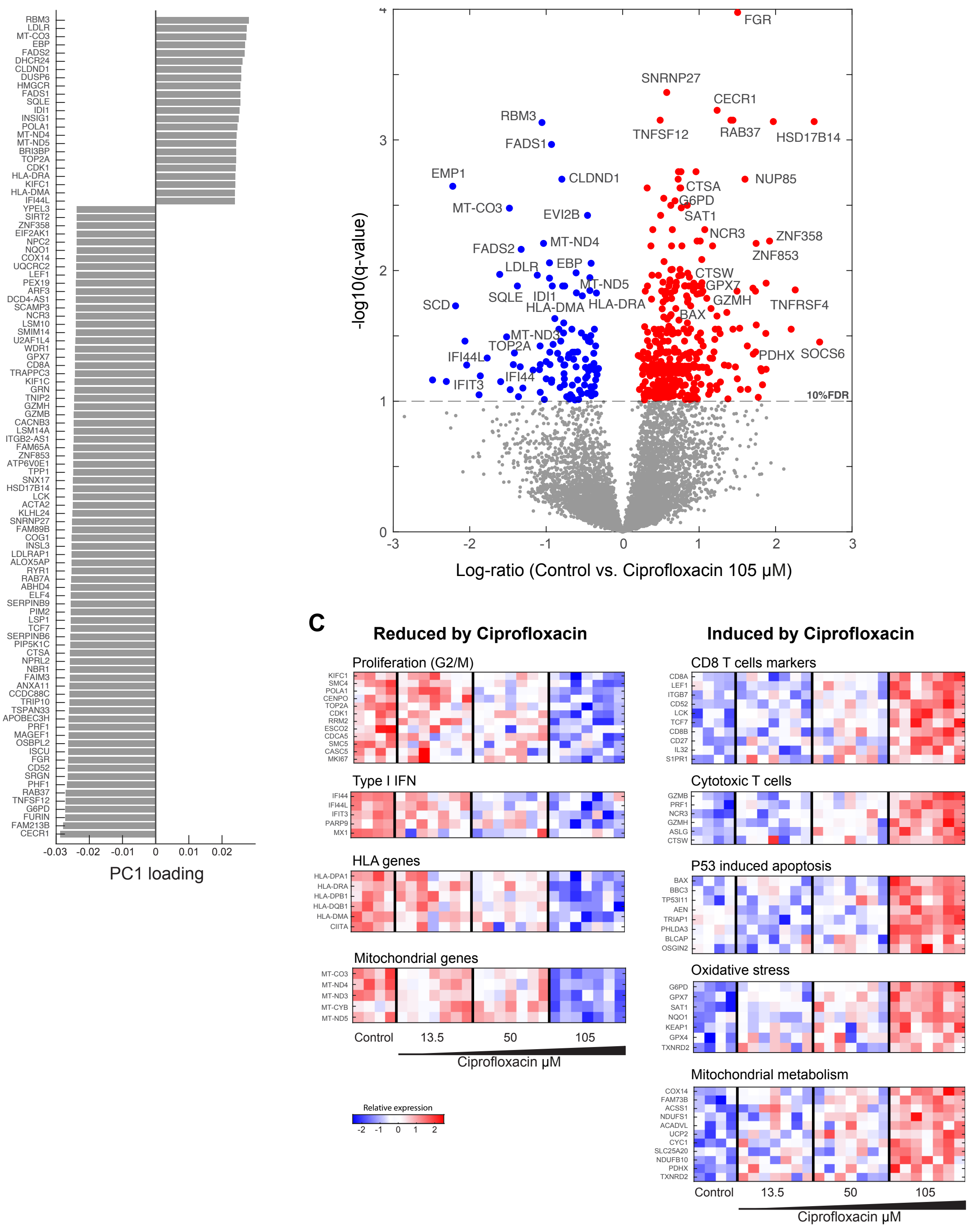


**Supplementary Figure 6: Dose-dependent transcriptomic shifts in proliferation and mitochondrial pathways in T cells following ciprofloxacin treatment.** T cells were activated from a single healthy PBMCs donor, then treated with the indicated concentrations of ciprofloxacin for 4 days and subjected to bulk RNAseq transcriptomics. (**A**) Shown are the top 100 genes with the highest PC1 loading. (**B**) Volcano plot of DEGs between untreated and ciprofloxacin (105 μM) treated samples (log fold change is presented on the X-axis, and -$\log_{10}$ q-value on the Y-axis). (**C**) Heatmaps of significantly DEGs (from B) grouped by biological function.

#### Supplementary Figure 7

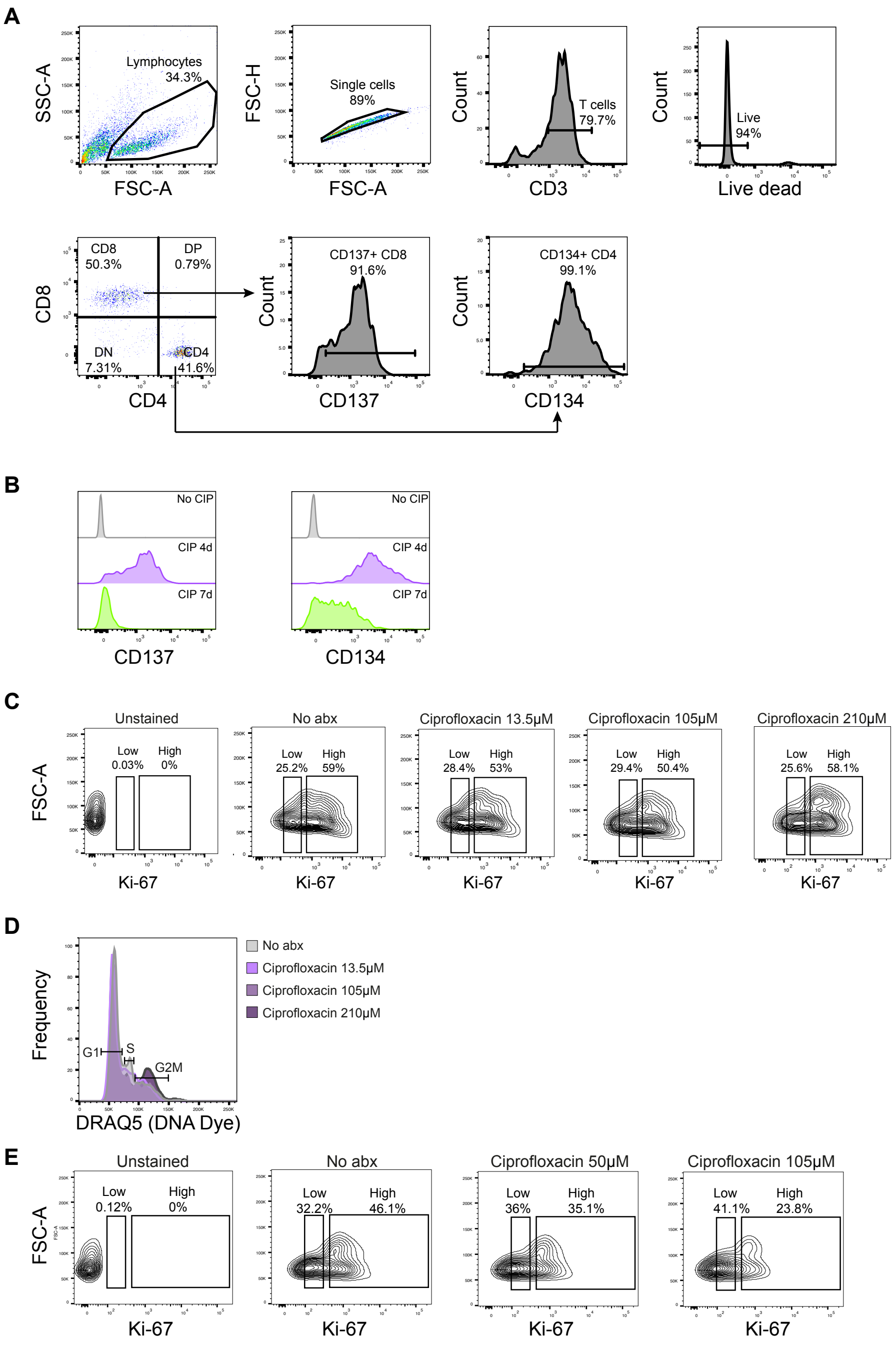


**Supplementary Figure 7: Ciprofloxacin inhibited T-cell proliferation.** (**A**) Gating strategy of T cell populations and activation markers. (**B**) Dynamics of T cell activation markers 4 days and 7 days after T cell expansion (unstained control in grey). CD137 is shown from the CD8 population, and CD134 is from the CD4 population. (**C**) T cells were treated with indicated concentrations of ciprofloxacin for 24h, stained with an antibody against Ki-67, and analyzed using flow cytometry. Gating for Ki-$67^{high}$ and Ki-$67^{low}$ is shown. (**D**) Ki-$67^{high}$ T cells from C were analyzed using DRAQ5 DNA-binding dye for analysis of cell cycle progression. **(E)** T cells were treated with the indicated concentrations of ciprofloxacin for 4 days, stained with an antibody against Ki-67, and analyzed by flow cytometry. abx- antibiotics. T cells were activated from PBMCs of a healthy donor.

#### Supplementary Figure 8

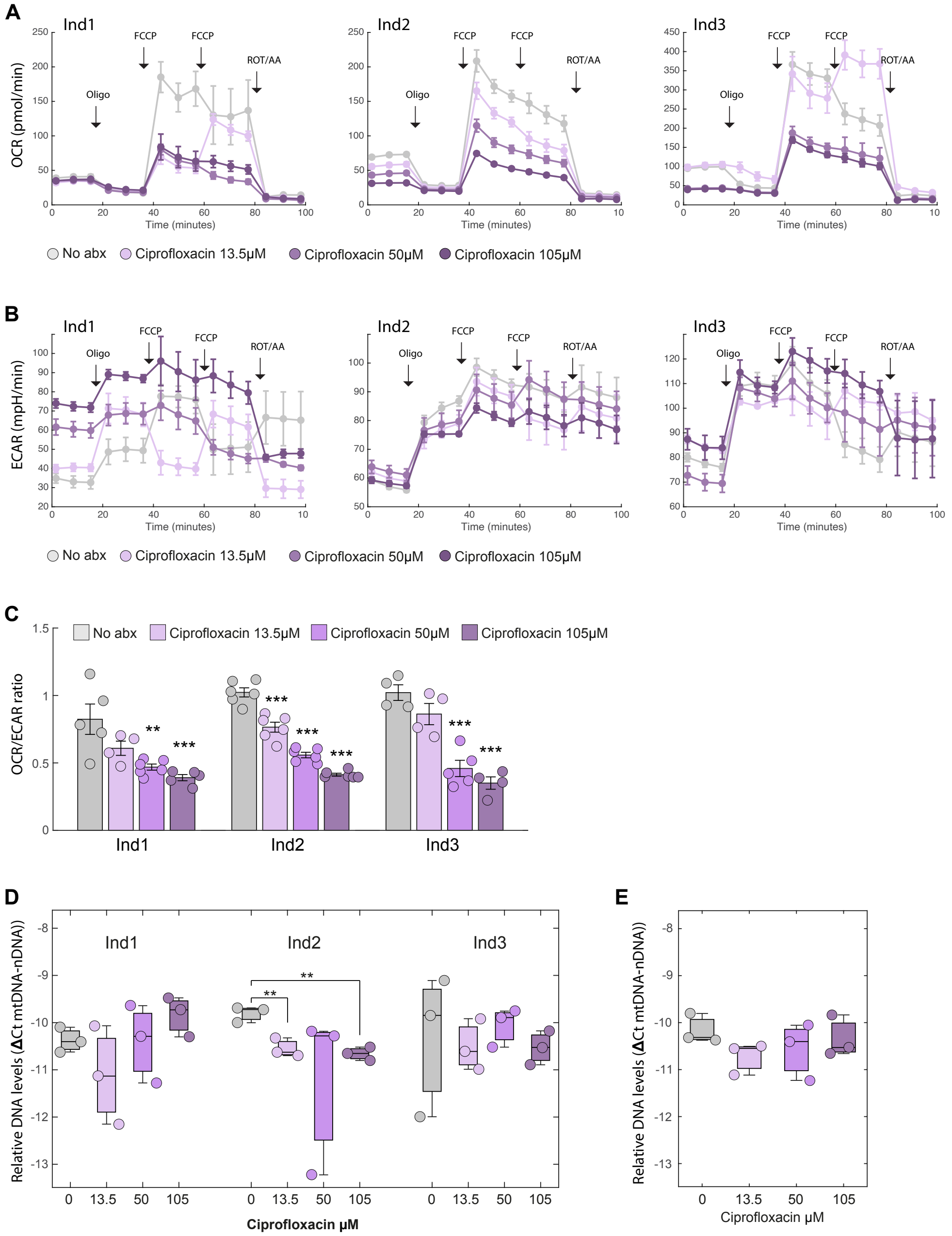


**Supplementary Figure 8: Dose-dependent impact of ciprofloxacin on mitochondrial respiration of T cells**. T cells from three healthy donors were treated with the indicated concentrations of ciprofloxacin or control for 4 days. Oxygen consumption rate (OCR) (**A**), extracellular acidification rate (ECAR) (**B**) were measured using a Seahorse XF extracellular flux analyzer. (**C**) OCR/ECAR ratio is shown for each donor per condition. Bars represent means ± SEM, dots represent technical replicates for each donor and condition. *P* values indicated are from one-way repeated-measures ANOVA followed by Dunnett's multiple comparison test, comparing each ciprofloxacin concentration with the untreated control (No abx). (**D-E**)

Relative mitochondrial DNA (mtDNA) to nuclear DNA (nDNA) ratio was measured from the same healthy donor samples, expressed as ΔCt (mtDNA - nDNA) for each individual (**D**) or averaged across the 3 individuals (**E**). *P* values indicated are from 2-sample t-test, relative to the untreated control (no antibiotics): ** $P < 0.01$, ***$P<0.001$.

### Supplementary Figure 9

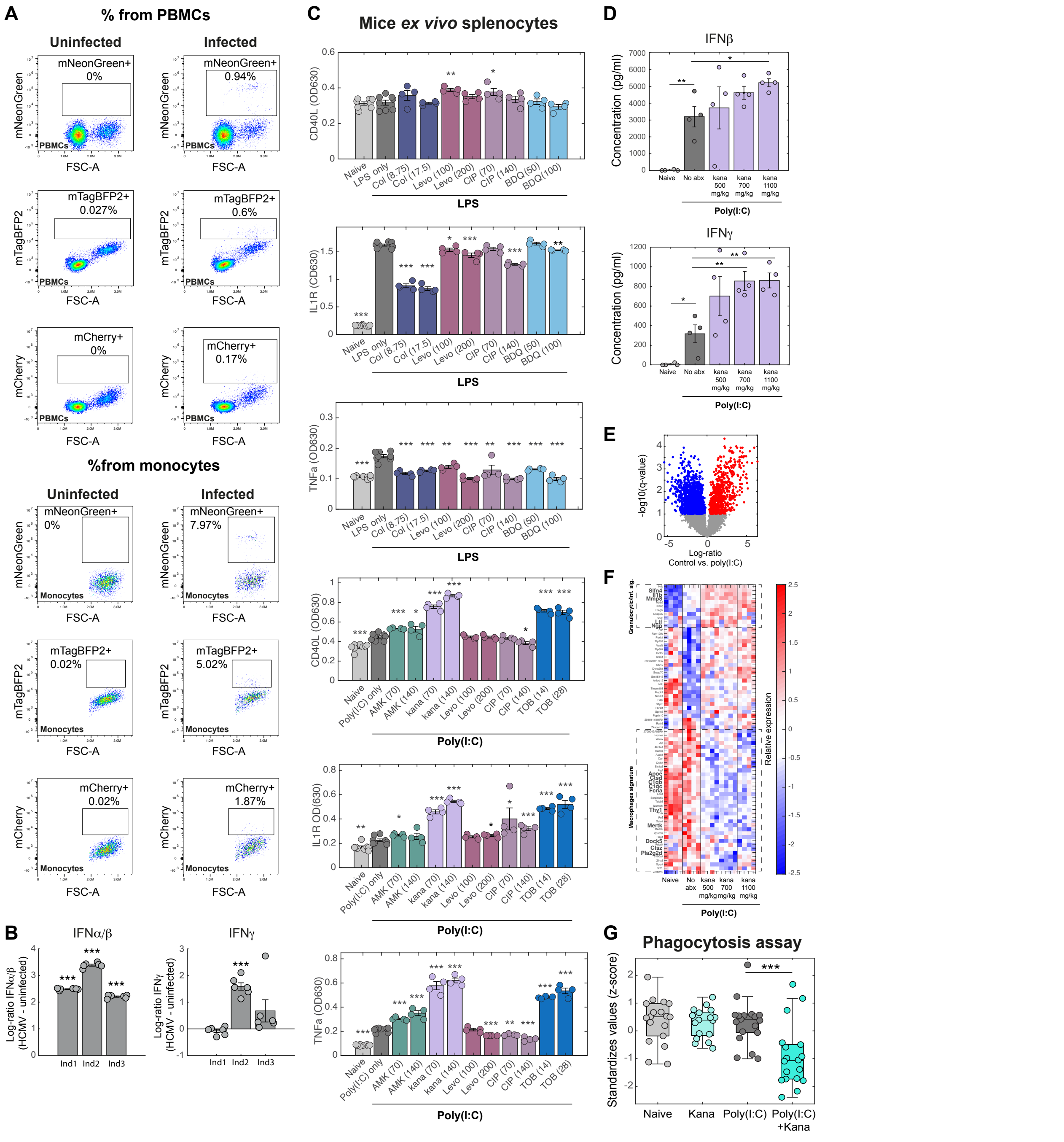


**Supplementary Figure 9: Immunomodulatory antibiotics affect cytokine secretion during viral infection.** (**A**) PBMCs from a healthy donor were infected with human cytomegalovirus (HCMV) carrying 3 fluorescent reporters: mNeonGreen under an immediate-early gene promoter, mTagBFP2 under an early gene promoter, and mCherry under a late gene promoter. Shown are flow cytometry plots of uninfected PBMCs and infected PBMCs 18 h post-infection, indicating infected cells based on the fluorescence signal of HCMV. The percentage of infected cells is shown from the total PBMCs and from the monocyte population, as indicated. Mono. - monocytes. (**B**) PBMCs from three healthy donors were infected with HCMV. After 18 h IFNα/β (left panel) and IFNγ (right panel) cytokine secretion was measured. Y-axis indicates the log-ratio of cytokine secretion of infected treatment vs. uninfected control, measured by HEK-blue reporter cell lines. (**C**) Splenocytes were extracted from naïve mice, exposed to the indicated stimulants ex vivo, and treated with the indicated antibiotics (X-axis: Col- colistin, AMK-amikacin, Kana-kanamycin, Levo-levofloxacin, CIP-ciprofloxacin, TOB-tobramycin, BDQ-bedaquiline) for 24h. The concentration (μg/ml) used per drug appears in parentheses. The measured cytokine levels appear on the Y-axis. (**D-F**) Mice were injected intravenously with poly(I:C) (150 μg/mouse), followed immediately by subcutaneous administration of kanamycin (500, 700, or 1,100 μg/kg). At 4 h post-injection, mice were sacrificed; blood was collected via retro-orbital bleeding for plasma cytokine analysis (**D**), and isolated splenocytes were subjected to RNA-seq (**E, F**). (**D**) Plasma levels of IFNβ (top panel) and IFNγ (bottom panel) quantified by ELISA. Each symbol represents an individual animal. (**E**) Volcano plot displaying differentially expressed genes (DEGs) between control and poly(I:C)-injected mice. Significantly upregulated and downregulated genes (10% FDR) are indicated in red and blue, respectively. (**F**) Heatmap of genes exhibiting significant differential expression (10% FDR) across kanamycin doses. Genes belonging to granulocyte inflammatory or macrophage-related signatures are highlighted in bold. (**G**) THP1 human monocytes were differentiated with PMA 100 ng/ml for 48h, followed by 48h resting and then exposed for 24h to supernatant from PBMCs that was collected from the following conditions: naive (no stimulation, no antibiotic), kana only, poly(I:C) only, poly(I:C) + kana. Latex fluorescent beads were added to THP1 cells for 3h, and phagocytosis was assessed by flow cytometry. (kana- kanamycin, 40 μg/ml). *P* values indicated are from a 2-sample t-test, * $p<0.05$, ** $p<0.01$, *** $p<0.001$.
